# GeneSegNet: a deep learning framework for cell segmentation by integrating gene expression and imaging

**DOI:** 10.1101/2022.12.13.520283

**Authors:** Yuxing Wang, Wenguan Wang, Dongfang Liu, Wenpin Hou, Tianfei Zhou, Zhicheng Ji

## Abstract

When analyzing data from in situ RNA detection technologies, cell segmentation is an essential step in identifying cell boundaries, assigning RNA reads to cells, and studying the gene expression and morphological features of cells. We developed a deep-learning-based method, GeneSegNet, that integrates both gene expression and imaging information to perform cell segmentation. GeneSegNet also employs a recursive training strategy to deal with noisy training labels. We show that GeneSegNet significantly improves cell segmentation performances over existing methods that either ignore gene expression information or underutilize imaging information.

## Background

RNA in situ hybridization (ISH) is a type of technology that measures the spatial locations of RNAs within tissue sections [1, 2, 3, 4, 5]. It provides great potential to understanding the spatial organizations of different cell types within a tissue, how different cell types interact with each other, and sizes and morphologies of cells. The RNA spatial location information is often accompanied by fluorescent staining (e.g., DAPI) images indicating locations of cells. However, the fluorescent images are noisy and cell boundaries are not directly captured and need to be inferred computationally. Accurately identifying cell boundaries is crucial for downstream cell-level analysis that relies on the identities and characteristics of individual cells.

There are two major categories of methods to identify cell boundaries. The first type of methods segment the image into small domains representing cell instances. They were originally designed for biomedical images without gene expression information and only utilize the imaging information. A classical example is the Watershed algorithm [6, 7], and methods based on deep neural network (DNN) techniques such as U-Net [8], Deepcell [9], and Cellpose [10] have also been developed in recent years. A fundamental limitation of these methods is that imaging information alone may not be enough to accurately capture cell boundaries. For example, these methods are more likely to capture cell nuclei boundaries instead of cell boundaries when only nuclei staining is available. Plus, their performances are affected by noise in images and training labels. The second type of methods utilize spatial locations of RNA reads to infer cell boundaries. These methods include Baysor [11] and JSTA [12]. Baysor assigns RNAs to individual cells and optionally accepts prior cell segmentation results obtained using the first type of methods such as Watershed. JSTA improves initial Watershed cell segmentation by iteratively reassigning boundary pixels based on cell type probabilities. A major limitation of these methods is that the imaging information is not directly used to obtain cell boundaries. As a result, cell boundaries do not fully match the actual images and can appear rather artificial in real applications. Plus, Baysor requires an estimate of cell sizes as input and may yield problematic results if the cell size estimate is biased. JSTA requires reference single-cell RNA-seq and cell type annotations as inputs which are not always available in real data. Another method SSAM [13] directly assigns RNAs to cell types and does not provide cell segmentation.

Cell segmentation methods based on DNN techniques have shown superior performances in extracting information from extensive amounts of images with annotated cell segmentation labels [14, 15]. In current practice, the training cell segmentation labels are assumed to be unambiguous and accurate. However, this assumption often does not hold. Accurately annotating cell masks is challenging even for experienced experts, due to difficulties such as low contrast of cell boundaries and cell adhesion. Moreover, manual annotations are subjective in general, and collecting cell segmentation annotations of high quality is time-consuming and expensive, making it almost prohibitive to manually clean or correct the labels in large-scale. Consequently, existing cell segmentation datasets tend to contain inaccurate labels (Fig. 1). Training DNNs on such noisy labeled datasets may cause suboptimal performance because it has been proved that DNNs easily overfit to noisy labels given their powerful learning ability [16]. Although the problem of learning from noisy labels has attracted growing attention in the field of deep learning [17, 18], this fundamental challenge is less touched by recent DNN-based cell segmentation methods.

**Figure 1.**
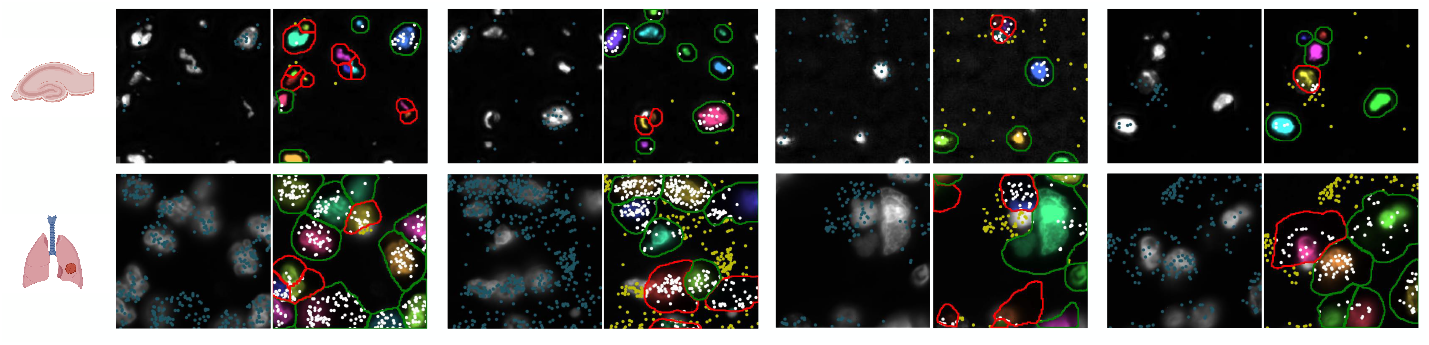
Examples of noisy training label annotations in hippocampus and NSCLC datasets. Four examples are given for each dataset. The left image in each example shows the image with detected RNAs (marked in blue). The right image indicates the corresponding annotations, where the potential noisy annotations are marked in red. RNAs falling inside annotated cell regions are marked in white and RNAs falling outside annotated cell regions are marked in yellow.

## Results

To address these issues, we propose GeneSegNet, a deep-learning-based cell segmentation method (Fig. 2). GeneSegNet integrates both RNA locations and imaging information within a single unified and fully differentiable network architecture, which enables both visually plausible and biologically reasonable cell segmentation. The rationale is that regions with high densities of RNAs are more likely to be within cells, and these regions may not have high pixel values in images such as nucleus staining. By converting discrete RNA spatial locations into a “pseudoimage” of 2D continuous probabilistic location maps, GeneSegNet accounts for the stochasticity of RNA spatial locations and both RNA and imaging information can be simultaneously fed into the neural network as inputs. This allows GeneSegNet to effectively extract and deeply fuse meaningful features from the two modalities throughout the whole neural network. To tackle the challenge of noisy training labels, GeneSegNet is trained through a recursive strategy: network predictions of the previous training round are refined and used as new training labels for the next-round network training. In this way, GeneSegNet boosts the robustness by making a joint optimization of learning network parameters and approximating true training labels, as opposed to previous methods treating the noisy labels as fixed. Such a recursive training strategy is able to correct inaccurate labels and hence improve the performance of GeneSegNet.

**Figure 2.**
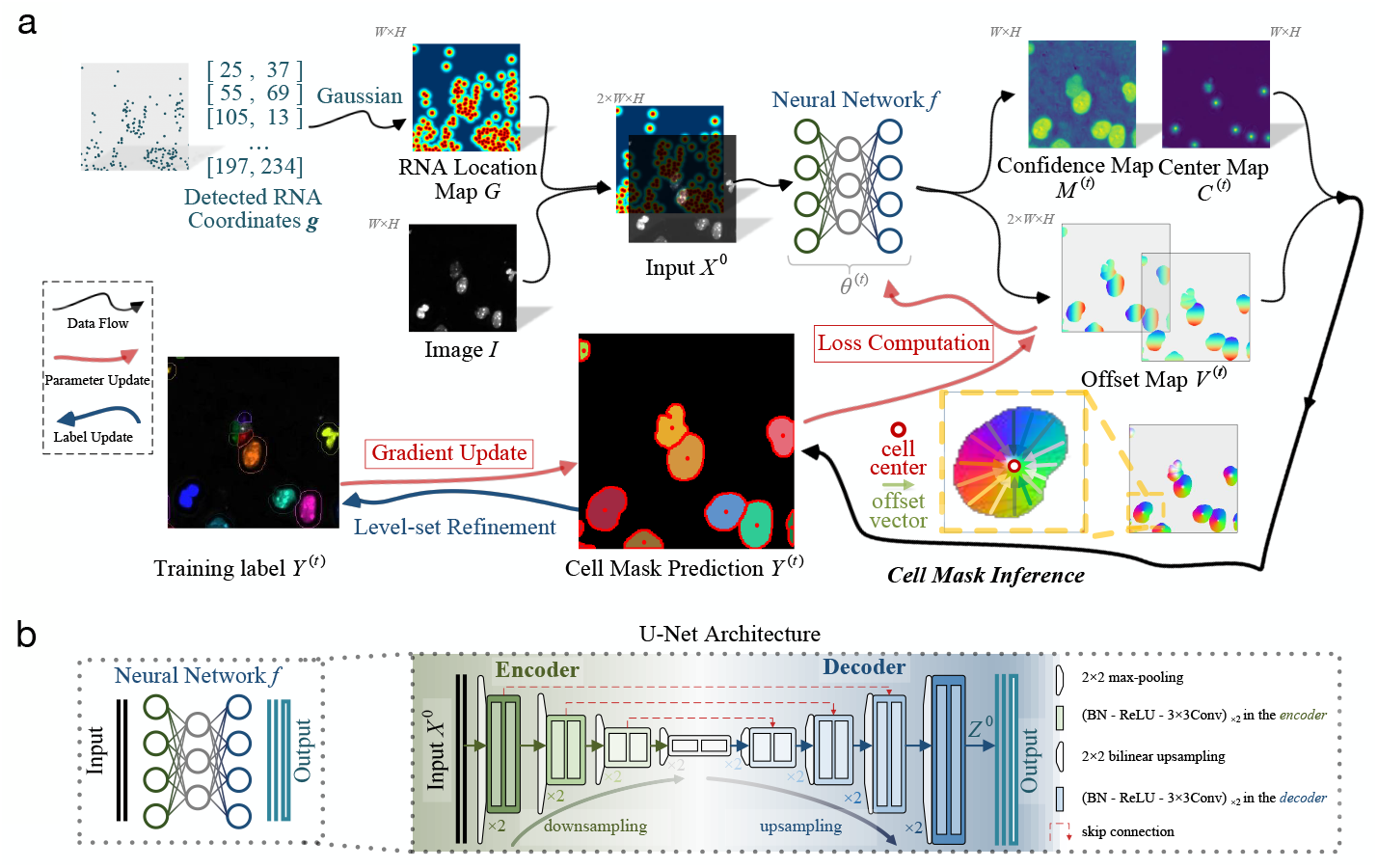
**a**, Overview of GeneSegNet framework. GeneSegNet makes a joint use of gene spatial coordinates and imaging information for cell segmentation, and is recursively learned by alternating between the optimization of network parameters and estimation of training labels for noise-tolerant training. The outputs of GeneSegNet include a *confidence map* 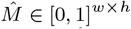 that stores the probability of each pixel being inside cell regions, a *center map Ĉ* ∈ [0, 1]^*w×h*^ that gives the likelihood of each pixel being a cell center, and a two-channel *offset map* 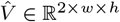 indicating the offset vector of each pixel to the center of its corresponding cell instance. Combining the outputs together, GeneSegNet can recover the precise boundary of each individual cell. **b**, GeneSegNet is built upon the general U-Net network architecture, where an encoder progressive downsamples the inputs for more expressive feature extraction, and a decoder upsamples the feature maps in a mirror-symmetric fashion for fine-grained cell segmentation.

To evaluate the performance of GeneSegNet, we applied GeneSegNet to a simulation dataset with ground truth information of cell boundaries and assignments of RNA spatial spots to cells (Methods), a real dataset of human non-small-cell lung cancer (NSCLC) generated by Nanostring CoxMx platform [19], and a real dataset of mouse hippocampal area CA1 (hippocampus) generated by in situ RNA detection [20]. The simulation dataset was designed to consider various scenarios of sparsely or densely distributed cells, high or low levels of image noise, and abundant or depleted amounts of genes. For real datasets, RNA reads are sparse in the hippocampus dataset and dense in the NSCLC dataset, representing diverse spatial patterns (Fig. 1). We compared GeneSegNet to two types of competing methods: image-based methods of watershed algorithm and Cellpose, and gene-based methods of Baysor and JSTA. Baysor was performed both with or without prior cell segmentation as input. For NSCLC dataset, both cell nucleus and membrane staining information are available. Cellpose applied to nucleus and membrane staining was used as the ground truth of cell segmentation. Cellpose applied to only nucleus staining was treated as a competing method, and GeneSegNet and Watershed algorithm were performed with only nucleus images as well. JSTA was only applied to real datasets since the cell type information, which is required as input by JSTA, was not generated for the simulation datasets.

Fig. 3, Fig. 4, and Fig. 5 show examples of cell images and RNA spatial locations, the ground truth of cell boundaries, and cell segmentation results by GeneSegNet and competing methods in simulation studies, NSCLS dataset, and hippocampus dataset, respectively. Compared to image-based methods of Watershed algorithm and Cellpose that do not consider RNA information, GeneSegNet generates larger cell boundaries which include more RNA reads within cells. Plus, GeneSetNet is less likely to oversegment cells. Gene-based methods of Baysor and JSTA perform poorly. Both methods tend to oversegment or miss entire cells in some cases. JSTA generates artificial circles and fails to include considerable amounts of RNA reads and some pixels with high pixel values within cell boundaries. Baysor uses a kernel density estimate (KDE) approach that is purely based on RNAs to obtain cell boundaries. The cell boundaries mismatch with actual images and result in unrealistic cell shapes. In summary, GeneSegNet benefits from both imaging and gene expression information, and generates smooth cell boundaries that capture actual cell shapes, include more RNA reads, and are less susceptible to imaging noise and oversegmentation.

**Figure 3.**
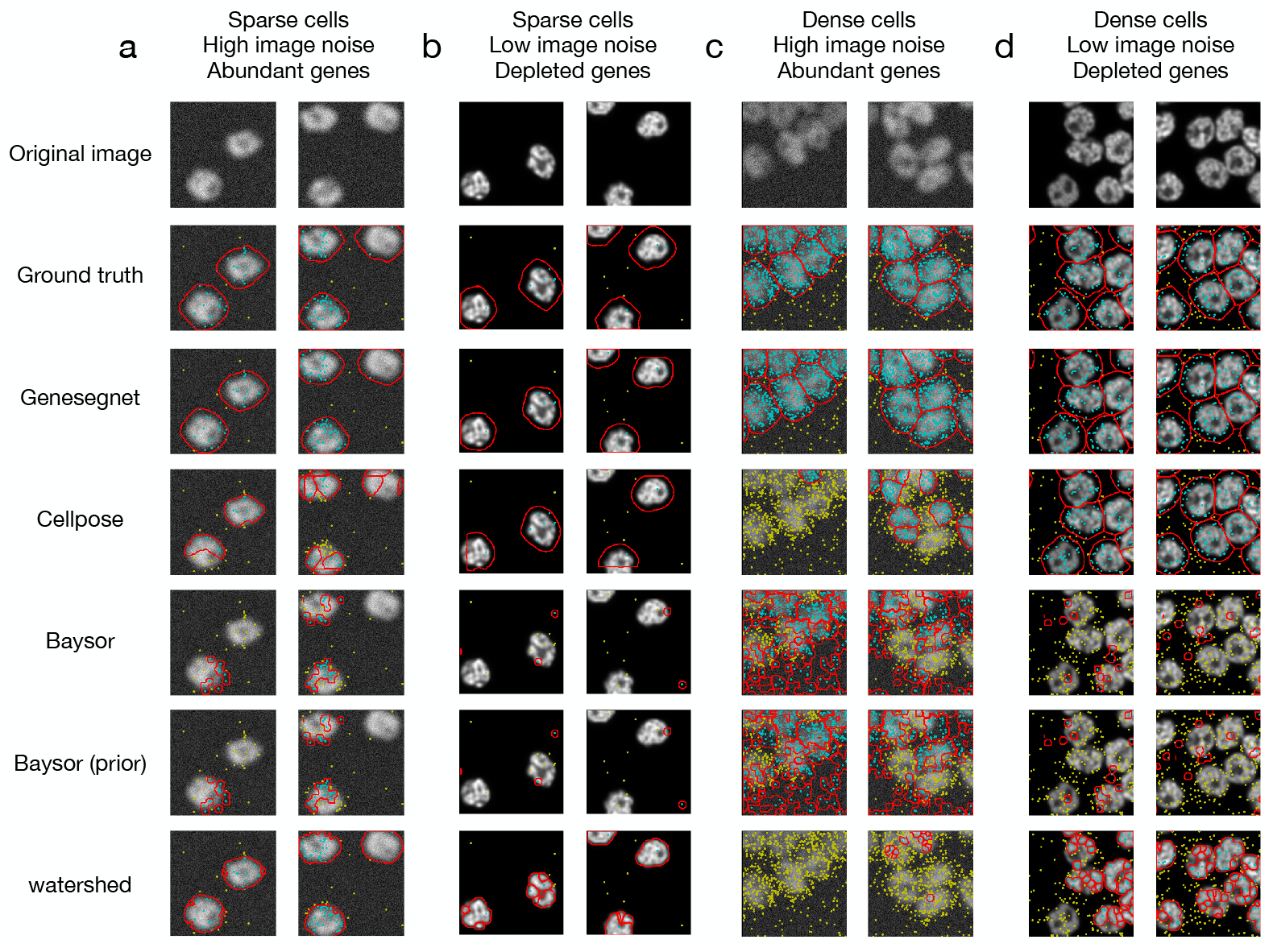
Examples of cell segmentation in the simulation dataset. The first row shows the original image and the second row shows the ground truth. Other rows show the results from cell segmentation methods. Each column is an example. The detected cell boundaries are marked in red. Light blue dots represent RNAs falling within cell boundaries. Yellow dots represent RNAs falling outside of cell boundaries. **a**, Sparsely distributed cells with high image noise and abundant amount of genes. **b**, Sparsely distributed cells with low image noise and depleted amount of genes. **c**, Densely distributed cells with high image noise and abundant amount of genes. **d**, Densely distributed cells with low image noise and depleted amount of genes.

**Figure 4.**
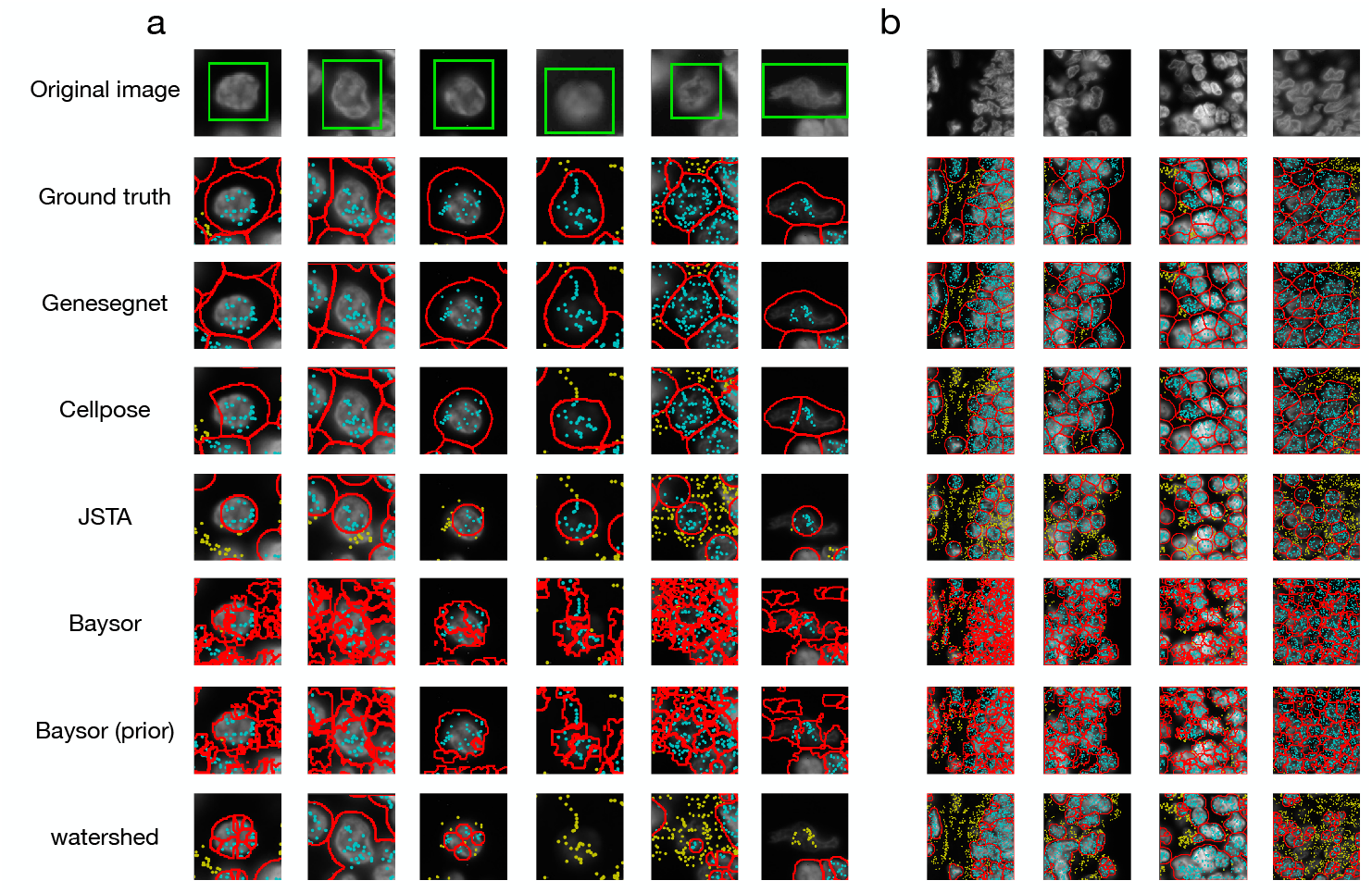
Examples of cell segmentation in the NSCLC dataset with zoomed in (**a**) and zoomed out (**b**) views. The first row shows the original image and the second row shows the ground truth. Other rows show the results from cell segmentation methods. Each column is an example. The detected cell boundaries are marked in red. Light blue dots represent RNAs falling within cell boundaries. Yellow dots represent RNAs falling outside of cell boundaries. In the zoomed in view (**a**), the cell of interest is highlighted in a green window.

**Figure 5.**
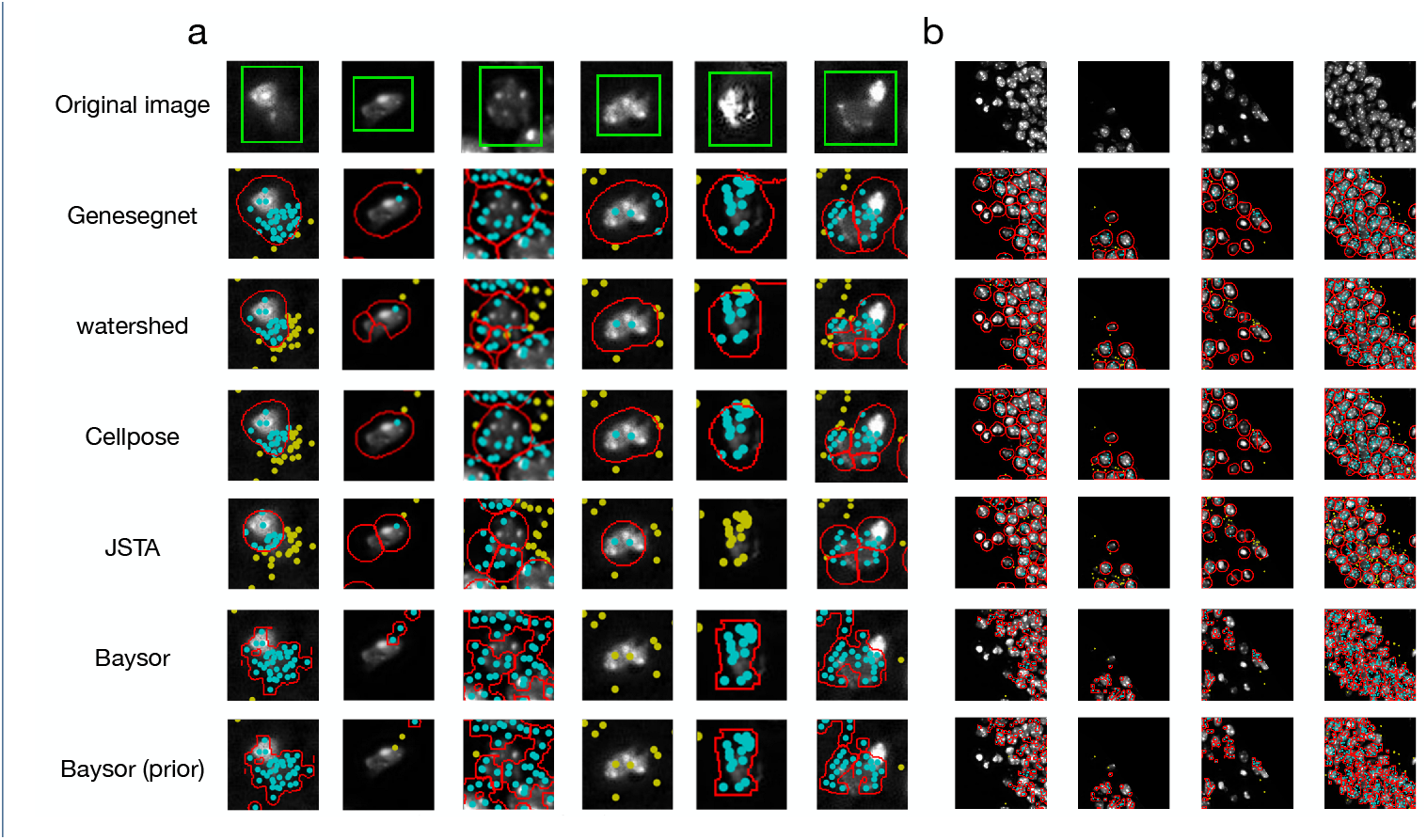
Examples of cell segmentation in the hippocampus dataset with zoomed in (**a**) and zoomed out (**b**) views. The first row shows the original image. Other rows show the results from cell segmentation methods. Each column is an example. The detected cell boundaries are marked in red. Light blue dots represent RNAs falling within cell boundaries. Yellow dots represent RNAs falling outside of cell boundaries. In the zoomed in view (**a**), the cell of interest is highlighted in a green window.

We next performed a global and systematic evaluation of all methods. We begin by computing the Intersection over Union (IoU) scores for images and genes in the simulation dataset, following protocols in existing studies [21, 22, 23, 8, 10, 24]. The IoU scores measure the similarity of cell boundaries (image IoU, Fig. 6a) and assignments of RNA spots to cells (gene IoU, Fig. 6b) between cell segmentation generated by GeneSegNet or a competing method and the ground truth. Methods with cell segmentation more similar to the ground truth have higher IoU scores. In the four simulation scenarios, GeneSegNet consistently outperforms other methods in both image IoU and gene IoU scores. Cellpose, which is also a deep-learning-based method but does not utilize information of RNA spatial locations, has comparable performance as GeneSegNet when there is low image noise and depleted amount of genes (Fig. 3b,d, Fig. 6a-b), but its performance significantly decreases when the image noise is high (Fig. 3a,c, Fig. 6a-b). In comparison, GeneSegNet still maintains high image and gene IoU scores even when there is a limited amount of image or RNA spatial information, demonstrating the advantage of utilizing both types of information. Watershed and Baysor have far worse performance (Fig. 6a-b), consistent with the observations in Fig. 3. We further studied the performance of GeneSegNet in a crowded region when cells are all neighboring to each other and the spatial locations of RNA spots form a continuum. The performance of GeneSegNet only decreases slightly when cells are crowded and densely distributed (Fig. 3c-d, Fig. 6a-b) compared to the performance when cells are sparsely distributed (Fig. 3a-b, Fig. 6a-b), showing that GeneSegNet is able to delineate cell boundaries and assign RNAs to cells relatively well in a crowded region.

**Figure 6.**
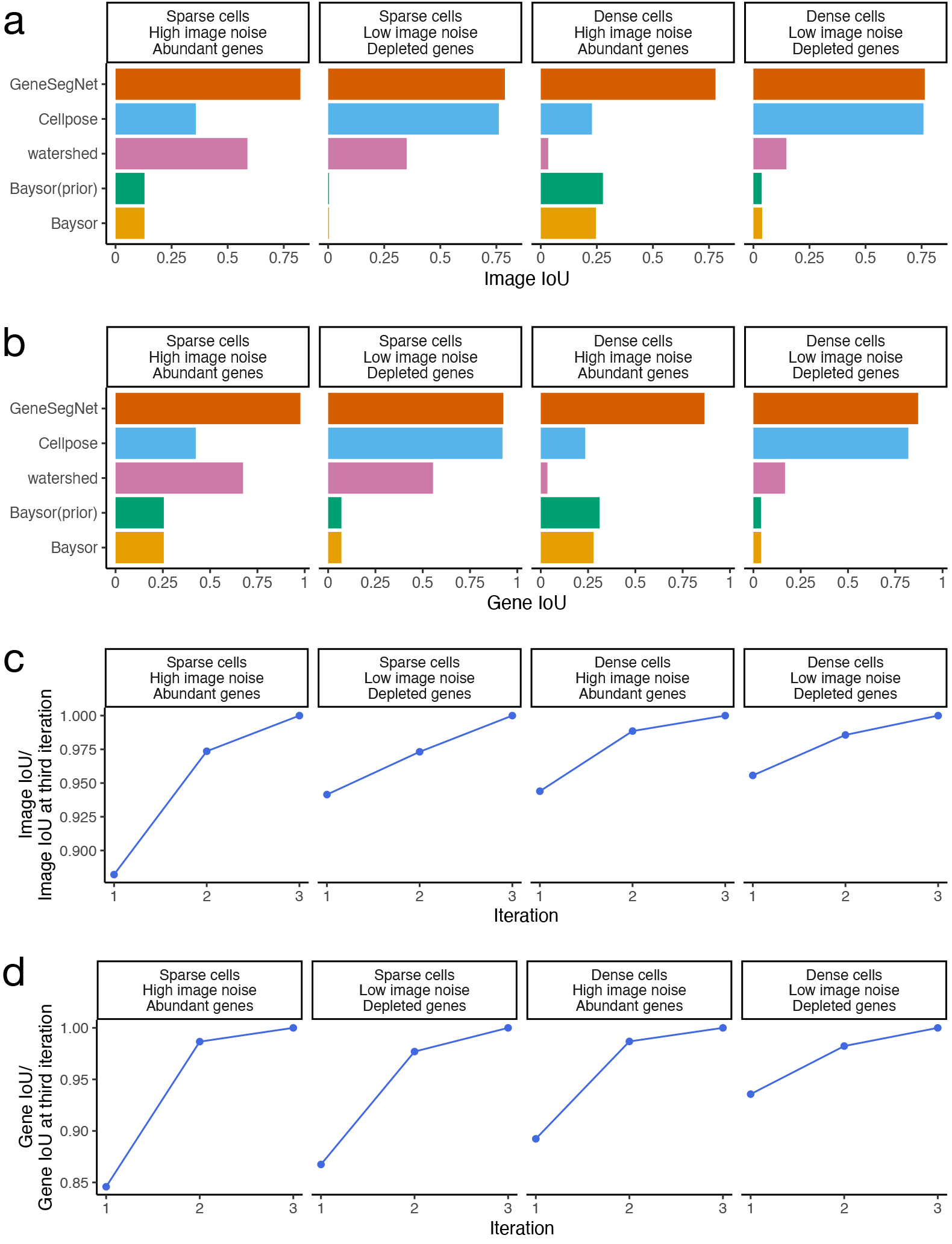
Quantitative analysis in the simulation dataset. **a**, The image IoU scores in four simulation scenarios. **b**, The gene IoU scores in four simulation scenarios. **c**, Image IoU score of the first three iterations divided by the image IoU score of the third iteration. **d**, Gene IoU score of the first three iterations divided by the gene IoU score of the third iteration.

To evaluate how the recursive training strategy employed by GeneSegNet contributes to improved cell segmentation, we also compared the gene and image IoU scores across the first three training iterations of GeneSegNet (Fig. 6c-d). Note that GeneSegNet with only the first training iteration is equivalent to the situation where the recursive training strategy is not applied. In all simulation scenarios, both image and gene IoU scores increase with the number of recursive training iterations, and using the recursive training strategy leads to 5% to 15% performance gains compared to not doing so. Moreover, the performance of the second and third iterations are highly similar, showing that the recursive training converges quickly and running three iterations is enough. Thus, GeneSegNet runs for three recursive training iterations by default.

We next evaluated the performance of cell segmentation methods on real datasets. Similar to the simulation study, we calculated gene and image IoU scores using the ground truth derived from the nucleus and membrane markers in the NSCLC dataset. GeneSegNet again outperforms other methods and has a significantly higher gene IoU compare to Cellpose (Fig. 7a-b), showing the improved ability of GeneSegNet to correctly assign RNA spots to cells. Since the hippocampus dataset does not have ground truth of cell boundaries, we then performed numerous analyses independent of the ground truth. We designed a cell calling score to test the ability of each method to absorb RNA reads near a cell into cell boundaries (Methods). GeneSegNet outperforms all methods in both real datasets (Fig. 7c and Fig. 8a). GeneSegNet and image-based methods have comparable performances to include pixels with high pixel values within cell boundaries, while gene-based methods fail to assign many high pixel values within cell boundaries (Fig. 7d and Fig. 8b). In comparison, GeneSegNet and gene-based methods have comparable performances to include RNAs residing in pixels with low pixels values within cell boundaries, while image-based methods have significantly lower performance (Fig. 7e and Fig. 8c). These results demonstrate that GeneSegNet is able to fully utilize both gene expression and imaging information, while existing methods only focus on a single information source. As a result, GeneSegNet is able to capture more cells with larger numbers of reads assigned to each cell (Fig. 7f and Fig. 8d) and larger cell sizes (Fig. 7g and Fig. 8e). Consistent with observations in Fig. 4 and 5, JSTA yields artificial cell boundaries with elongation close to 1 (representing a circle), while other methods yield more realistic cell boundary elongations (Fig. 7h and Fig. 8f). Baysor segmentation also yields significantly lower convexity compared to other methods (Fig. 7i and Fig. 8g), which is unrealistic in many cases and is consistent with observations in Fig. 3, Fig. 4, and Fig. 5.

**Figure 7.**
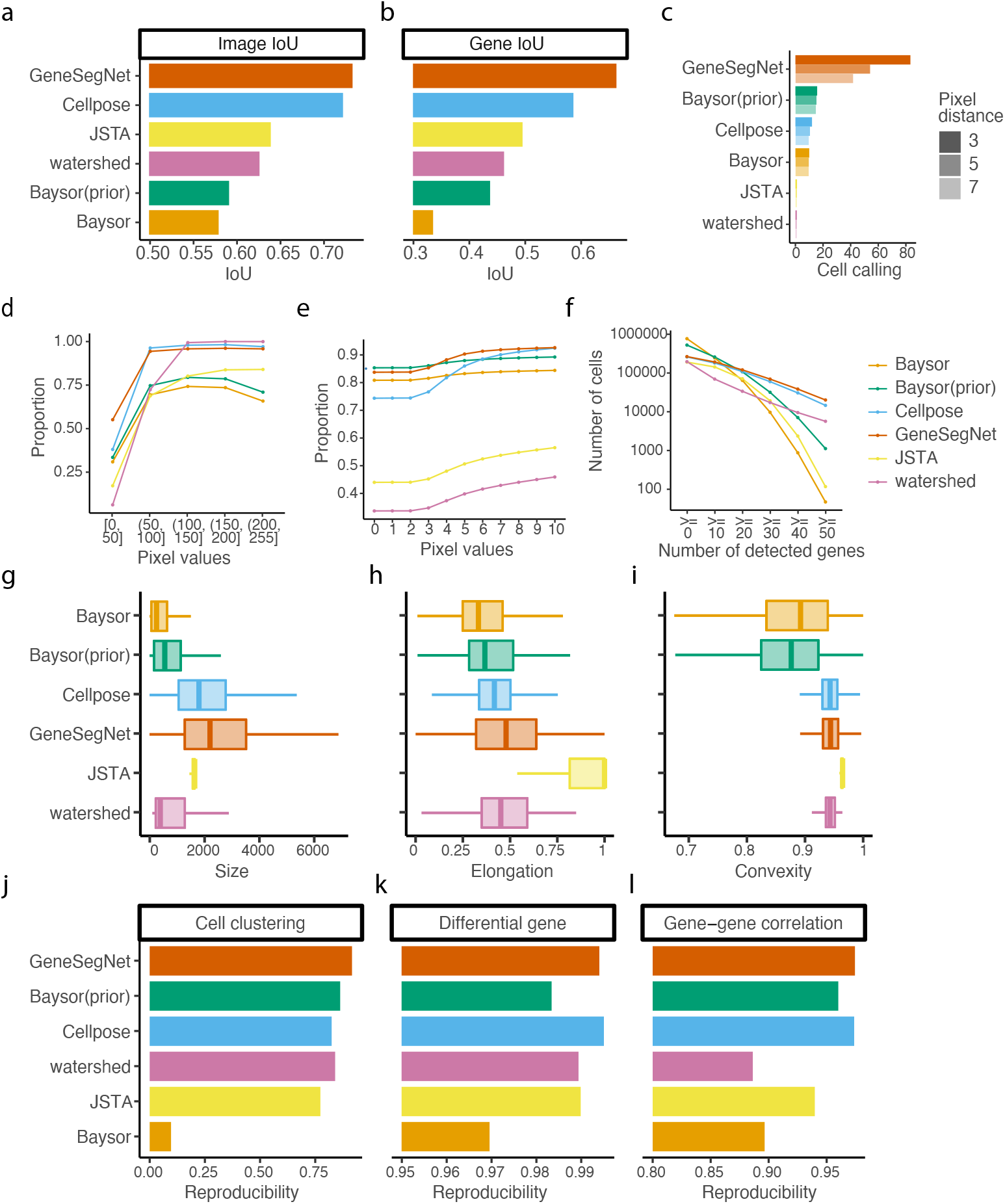
Quantitative analysis in NSCLC dataset. **a**, The image IoU scores. **b**, The gene IoU scores. **c**, Cell calling metrics. **d**, Proportion of pixels within cell boundaries (y-axis) for pixels with different levels of pixel values (x-axis). **e**, Proportion of RNA reads within cell boundaries (y-axis) for RNA reads with different pixel values (x-axis). **f**, Number of cells (y-axis) with different numbers of detected genes (x-axis). A gene is detected if it has at least one read in that cell. **g-i**, The distribution of cell size in pixels (**g**), elongation (**h**), and convexity (**i**). **j-l**, Reproducibility of cell clustering (**j**), differential gene analysis (**k**), and gene-gene correlations (**l**) across biological replicates.

**Figure 8.**
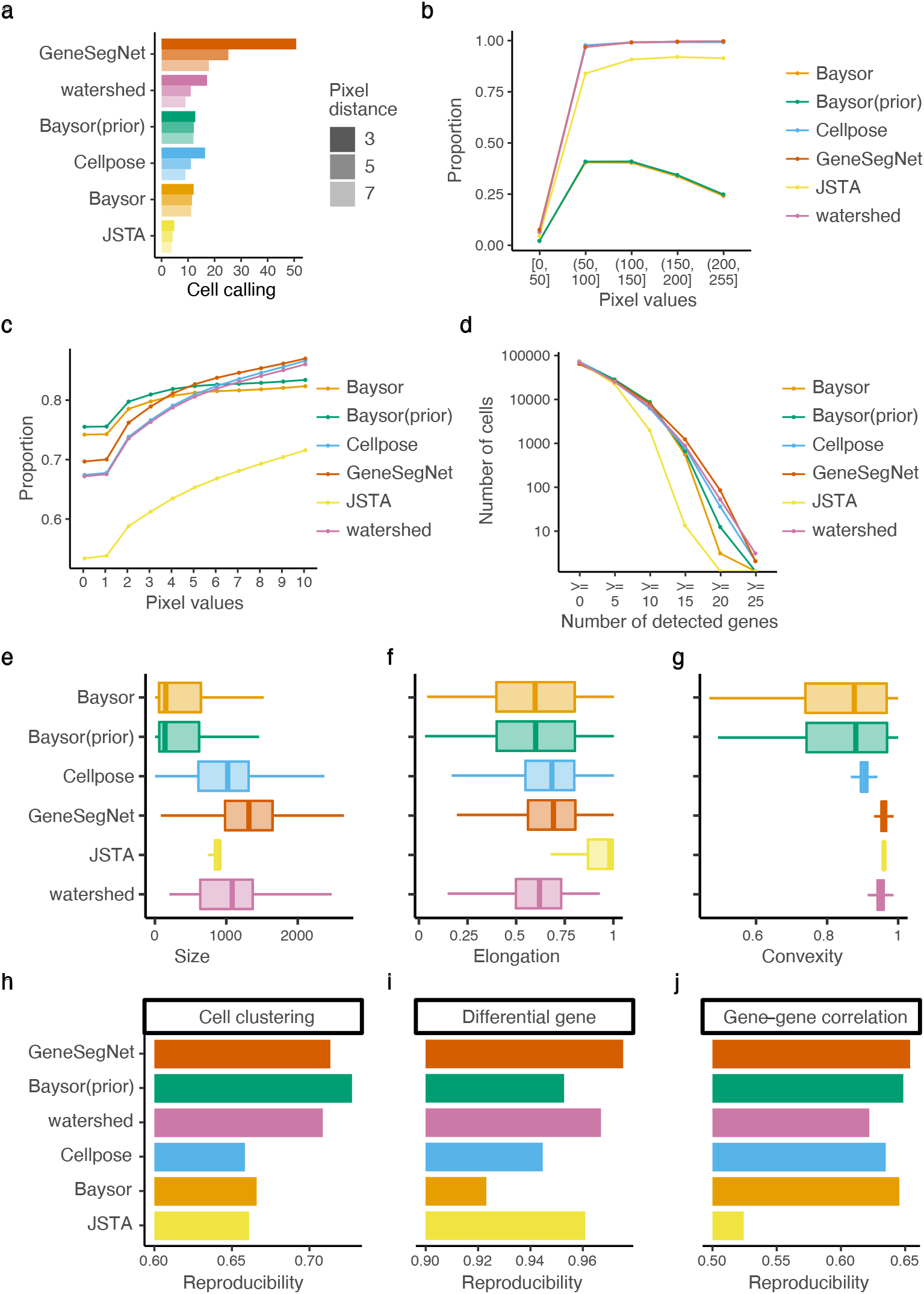
Quantitative analysis in hippocampus dataset. **a**, Cell calling metrics. **b**, Proportion of pixels within cell boundaries (y-axis) for pixels with different levels of pixel values (x-axis). **c**, Proportion of RNA reads within cell boundaries (y-axis) for RNA reads with different pixel values (x-axis). **d**, Number of cells (y-axis) with different numbers of detected genes (x-axis). A gene is detected if it has at least one read in that cell. **e-g**, The distribution of cell size in pixels (**e**), elongation (**f**), and convexity (**g**). **h-j**, Reproducibility of cell clustering (**h**), differential gene analysis (**i**), and gene-gene correlations (**j**) across biological replicates.

Finally, to demonstrate the impact of different cell segmentation methods on down-stream analysis, we performed cell clustering, differential gene analysis, and gene-gene correlation analysis using gene expression matrices created by RNA assignments to cells (Methods). Cell clustering groups cells with similar gene expression profiles into the same clusters and is useful to identify cell types of individual cells. Differential gene analysis identifies genes with differential gene expression levels across cell clusters that may drive the heterogeneity of cells. Gene-gene correlation identifies pathways of genes with similar expression patterns across cells that may share common biological functions, and can be used to construct gene co-expression networks for screening candidate biomarkers or therapeutic targets [25]. Since there is no gold standard in real datasets to directly assess these down-stream analysis, we tested if the results of cell clustering, differential gene analysis, and gene-gene correlations are reproducible across biological replicates (Methods), similar to existing studies [26, 27]. GeneSegNet again has the best overall performance in both datasets and for three types of reproducibility (Fig. 7j-l and Fig. 8h-j). Other competing methods perform poorly in at least one scenario. For example, Cellpose has significantly lower reproducibility compared to GeneSegNet for cell clustering and differntial gene analysis in the hippocampus dataset (Fig. 8h-i). These results demonstrate that down-stream analyses with GeneSegNet are more stable and reproducible.

## Discussion

Similar to other deep-learning-based cell segmentation methods, GeneSegNet cannot directly analyze a large image with high resolution as a whole due to the limitations of GPU memory. To deal with such large images, GeneSegNet follows a common approach and crops the large image into patches of smaller images, performs segmentation on the smaller images, and stitches results back to produce segmentation on the original large image. This approach may result in minor performance loss on the edges of the smaller images.

In this study, GeneSegNet and Cellpose were trained and tested on the same datasets with different data splits for evaluation purpose. In real practice, cell segmentation methods will be more user-friendly if a model pre-trained on one dataset can be directly applied to another new dataset without additional training. We further tested the performance of GeneSegNet and Cellpose when they are trained on one dataset and tested on a different dataset, and the gene and image IoU scores of GeneSegNet only decrease slightly (Additional file 1: Fig. S1). The degree of performance loss is slightly less than that of Cellpose, showing that GeneSegNet models have higher generalizability. Thus, a pre-trained model by GeneSegNet can be readily applied to a new dataset.

In addition to evaluations based on ground truth, this study also uses numerous metrics independent of ground truth, such as cell calling, proportion of pixels with high pixel values or with RNA reads that fall within cell boundaries, cell size and elongation, and reproducibility analysis. Note that these metrics cannot fully reflect the actual performance of the methods due to the lack of ground truth and their inherent limitations. For example, the ability to absorb more RNA spots into cell boundaries is only desirable when those RNA spots actually belong to the cells. A larger cell size by a segmentation method is only desirable if it reflects the actual cell size. Although these results serve as a good complement, they need to be interpreted jointly with the evaluations based on ground truth to comprehensively benchmark the performances of different cell segmentation methods. In addition, the ground truth for NSCLC dataset in this study is computationally derived from Cellpose. Although this ground truth is generally reliable (Additional file 2: Fig. S2), it may not exactly reflect the real cell boundaries and should be interpreted with care Although GeneSegNet’s recursive training strategy will lead to increased running time, GeneSegNet is still able to finish running within 24 hours for all the datasets with 8GB GPU memory (Additional file 3: Table S1). We expect the running time can be further decreased if a more powerful GPU is available.

GeneSegNet is a general framework. Its components, such as the U-Net architecture, can be replaced with more advanced machine learning architectures to appear in the future to further improve its performance. GeneSegNet also has the potential to be applied to other types of imaging data where information such as chromatin accessibility is simultaneously measured. GeneSegNet supports multiple image channels as input, and also accepts probability maps for cell segmentation as input, as long as the probability maps have the same data format as ordinary images.

## Conclusions

We present GeneSegNet, a deep-learning-based framework to perform cell segmentation for RNA in situ hybridization data. While existing cell segmentation methods predominantly rely on one source of information, GeneSegNet integrates RNA spatial information and imaging information under a unified U-Net architecture. In addition, GeneSegNet adaptively updates and optimizes the noisy training labels using a recursive training strategy. We show that GeneSegNet outperforms existing cell segmentation methods and improves the performance of downstream analysis.

## Methods

### Methods Overview

#### Overall Model

GeneSegNet exploits both imaging information and spatial locations of RNA reads for cell segmentation, based on a general U-Net architecture [8] (Fig. 2). Specifically, our GeneSegNet network *f*, parameterized by *θ*, takes as inputs an *intensity image I* ∈ [0, 1]^*W×H*^ and an *RNA location map G* ∈ [0, 1]^*W×H*^, both with *W ×H* spatial resolution. The location map *G* is constructed by putting 2D Gaussian kernels on the detected RNA positions of image *I*, for easing the encoding of RNA location information. The outputs of GeneSegNet include i) a *confidence map* 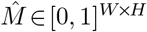 that stores the probability of each pixel being inside cell regions, ii) a *center map Ĉ* ∈ [0, 1]^*W×H*^ that gives the likelihood of each pixel being a cell center, and iii) a two-channel *offset map* 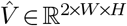 indicating the offset vector of each pixel to the center of its corresponding cell instance. Through the confidence map 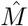, center map *Ĉ*, and offset map 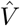, GeneSegNet can discover individual cells and recover their precise boundaries.

#### Recursive Training with Noisy Annotations

To address the issue of learning with ambiguous and noisy annotations, GeneSegNet is trained in a *recursive* fashion, *i*.*e*., alternatively optimizing network parameters and estimating true cell segmentation labels on training data. Specifically, given a set of *N* training images 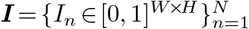, and corresponding RNA location maps 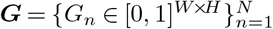, we denote the initial network parameters as *θ*^(0)^, and the initial labels for the confidence map, center map, and offset map as 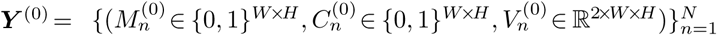, which are derived from the noisy cell mask annotations of training datasets (*cf*. §Network Training). The update rules of network parameters *θ* and labels ***Y*** can be described as:

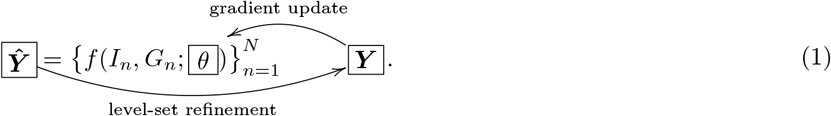

Specifically, at each recursive training round *t* = {0, 1, 2, *…* }, we have:

- *Update θ with fixed* ***Y*** : The network parameters *θ* are updated by the AdamW [28] optimizer on a cell segmentation loss function ℒ, which evaluates the network predictions ***ŷ***^(*t*)^ against current training labels ***Y***^(*t*)^:

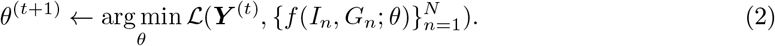

The detailed definition of the training loss *ℒ* is given in §Network Training.
- *Update* ***Y*** *with fixed θ*: We use the updated network parameters *θ*^(*t*+1)^ to make predictions ***ŷ***^(*t*+1)^ over the training data (***I, G***). Then the level-set algorithm [29] is used for boundary refinement. Level-set has been widely used in image segmentation applications, due to its decent boundary detection capacity, which can robustly represent cell contours with weak edges and inhomogeneities [30]. Then the training labels are updated as:

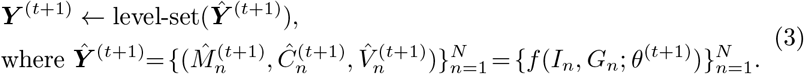

#### Cell Mask Inference

When performing cell mask inference, GeneSegNet leverages its outputs of confidence map 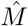, center map *Ĉ*, and offset map 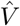, to group pixels into individual cell instances. Specifically, we first locate a set of cell centers {*ô*_1_, *ô*_2_, *…, ô*_*K*_} using center map *Ĉ*, where 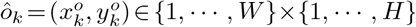 and *Ĉ*(*ô*_*k*_) *≥* 0.5. Next, we assign each pixel *p* = (*x, y*) ∈ {1, *…, W* } *×*{1, *…, H*} that falls in cell regions 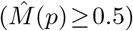 to *k*^*\**^-*th* cell center, according to:

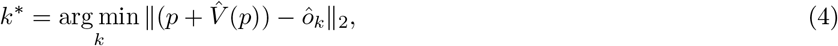

where 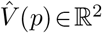 gives the center-pointing vector along *x*- and *y*-axes, *i*.*e*., 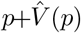 is expected to be the cell center of *p*. Hence the cell pixel *p* is clustered to the detected center *ô*_*k*_*\** that is nearest to 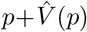.

All notations are explained in Additional file 4: Table S2. Next, we articulate each of the contributing components of this study.

### Datasets

#### Mouse hippocampus CA1 region (hippocampus) dataset

In this dataset, in situ sequencing technology was used to identify the spatial locations of RNA transcripts in 28 samples of mouse hippocampus CA1 regions [20]. Each sample has spatial locations of RNA reads and a one-channel grayscale image stained by DAPI. Watershed segmentation was conducted by the original study based on the DAPI stain. The image resolution varies from 3560*×*3034 to 6639*×*5548 pixels. The spatial locations of RNA reads and the DAPI stain images were used as inputs to GeneSegNet, and the watershed segmentation was used as the initial cell instance labels for GeneSegNet model training. The spatial locations of RNA reads, DAPI images, and watershed segmentation were directly downloaded from the original publication [31].

#### Human non-small cell lung cancer (NSCLC) dataset

In this dataset, NanoString CosMx™ platform was used to detect the spatial locations of RNA in human NSCLC tissues [19]. Two imaging samples from the same NSCLC tissue were used in this study. Each image sample provides 5 stains with both nuclear and membrane markers (Membrane stain, PanCK, CD45, CD3, DAPI), and each image is a one-channel grayscale image. Cellpose [10] was used to perform cell segmentation using both nuclear and membrane images by the original study. Each NSCLC sample contains around 30 high-resolution images with RNA reads information and the image resolution is 5472 × 3648 pixels. The spatial locations of RNA reads and the DAPI stain images were used as inputs to GeneSegNet, and the Cellpose segmentation was used as the initial cell instance labels for GeneSegNet model training. The spatial locations of RNA reads, DAPI images, and Cellpose segmentation were directly downloaded from NanoString website [32].

#### Simulation dataset

We consider several scenarios in the simulation dataset: scenarios with images of high or low noise levels, scenarios with densely or sparsely distributed cells, and scenarios with abundant or depleted amount of genes. A specialized toolbox for generating digital phantoms of cell nuclei [33] was employed to simulate nucleus image data with ground truth of nuclei boundaries. For each simulation scenario, we generated 24 small images of size 1206 × 906 and then stitched the images into a whole image with a resolution of 7248 × 3624 pixels. The ground truth nuclei boundaries were stitched in the same way. To simulate cell boundaries, we used the `binary_dilation` function in `scipy` Python package with default parameters to perform a dilation process on each cell nuclei. This process was repeated 20 times to expand 20 pixels out of the boundary of each nuclei to generate its cell boundary. To simulate images with high noise level, “dynamic range usage” was set as 1% and “acquisition time” was set as 10000 *ms*. To simulate images with low noise level, “dynamic range usage” was set as 25% and “acquisition time” was set as 5000 *ms*. To simulate images with densely distributed cells, the “Amount” parameter was set as 99. To simulate images with sparsely distributed cells, the “Amount” parameter was set as 10. Volume of interest was set as 160 × 120 × 12 in all simulation scenarios. All other parameters were set as default values. In total, we simulated 264 cells for the scenario of sparsely distributed cells with high image noise, 264 cells for the scenario of sparsely distributed cells with low image noise, 2348 cells for the scenario of densely distributed cells with high image noise, and 2343 cells for the scenario of densely distributed cells with low image noise.

As the toolbox can only simulate images, we then employed a customized procedure to simulate spatial locations of RNA spots on top of the simulated images, using the real NSCLC dataset [19] as a source. In addition to the spatial locations of RNA spots, we also simulated gene assignments for RNA molecules, which are required by Baysor as input information. We first simulated a situation with abundant amount of genes. To simulate RNA spots within cells, for each simulated cell, we randomly selected one real cell from the NSCLC dataset. We then randomly selected pixels within the boundary of the simulated cell to be RNA spots and assigned genes to the RNA spots, so that the number of RNA molecules for each gene is the same as that of the selected NSCLC cell. To simulate RNA spots outside of cells, we calculated the ratio between the number of cells in the simulation dataset and the real NSCLC dataset. This ratio was multiplied by the total number of RNA molecules outside of cells in the NSCLC dataset to obtain the total number of RNA molecules outside of cells in the simulation dataset. We then randomly selected the desired number of RNA molecules as well as their gene assignments from the NSCLC dataset, and randomly assigned the RNA molecules to pixel outside cells in the simulation dataset. We next simulated a situation with depleted amount of genes by randomly subsampling 10% of all RNA molecules (regardless of within or outside of cells) from a situation with abundant amount of genes.

The simulated spatial locations of RNA molecules and simulated images were used as inputs to GeneSegNet. The initial cell instance labels for training GeneSegNet were obtained from the segmentation generated by a GeneSegNet model trained on the hippocampus dataset.

### Neural Network Architecture

#### Network Input

Driven by the premise that comprehensive use of imaging information and gene expression can facilitate cell segmentation, we intend to feed both the image *I* ∈ [0, 1]^*W×H*^ and RNA spatial positions 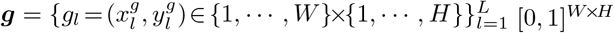 into our GeneSegNet network. To facilitate the simultaneous encoding of imaging and RNA information, we transform discrete RNA positions ***g*** into a single, continuous RNA location map *G* ∈ [0, 1]^*W×H*^, following the common practice in human pose estimation [34]. Specifically, the RNA location map is achieved by placing an unnormalized 2D Gaussian with a small variance *σ* (e.g., 7, 9) at each gene position.

Formally, the value of each position *p* =(*x, y*) in *G* is computed as:

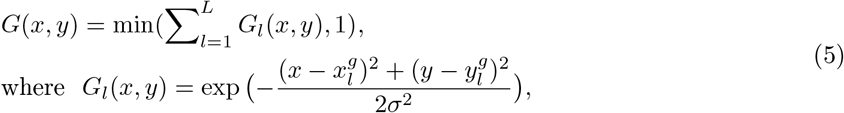

Here *G*_*l*_(*x, y*) encodes the confidence that a gene appears in coordinate (*x, y*), based on the *l*-th detected gene location, *i*.*e*., 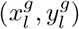. The RNA location heatmap *G* is an aggregation of the Gaussian peaks of all the *L* gene location distributions, *i*.*e*., 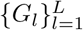, in a single “heatmap”, and we truncate each of its element to the range [0, 1] for numerical stability.

Based on the unified representation format of imaging information and RNA location information, the input to our GeneSegNet network is a two-channel “image” with *W ×H* spatial resolution:

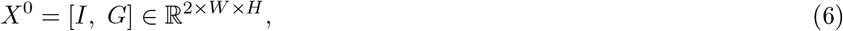

where [*·, ·*] denotes the matrix concatenation along the channel dimension. For *X*^0^, the first channel corresponds to the 2D density image *I*, and the second channel corresponds to the continuous RNA location map *G*.

#### Detailed Architecture

GeneSegNet is built upon the U-Net architecture [8, 10] (Fig. 2b), which is arguably the standard neural network architecture for medical image segmentation. Basically, U-Net downsamples convolutional features several times and then reversely upsamples them in a mirror-symmetric manner. This U-shaped architecture is implemented as a fully convolutional encoder-decoder network, which is composed of an *encoder* for downsampling/extracting the feature maps and a *decoder* for upsampling/reconstructing the feature maps. Specifically, the encoder conducts feature extraction at four spatial scales. At each scale, 2 × 2 max-pooling is first adopted to downsample the input feature map, followed by two residual blocks with additive identity mapping [35]. Each encoder residual block has two 3 × 3 convolutions, each of which is preceded by batch normalization (BN) and ReLU activation. Formally, the following operations are performed at *i*-*th* encoding scale:

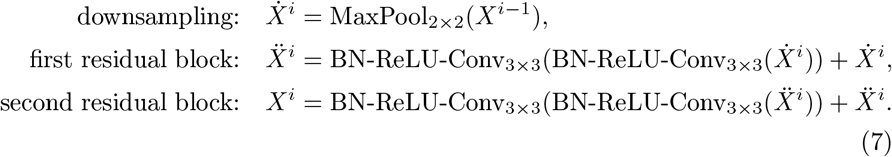

As such, the encoder repeatedly extracts more contextual feature maps *X*^*i*^ from preceding shallow feature maps *X*^*i-*1^, yet at the cost of reduced spatial resolution. The valuable information from both the image *I* and the RNA location map *G* can be automatically gathered and deeply fused.

To rebuild the spatial context for fine-grained segmentation, the decoder progressively upsamples the final most expressive yet coarse-grained feature maps of the encoder. More specifically, analogous to the encoder, the decoder conducts feature reconstruction at four spatial scales. At each scale, 2 × 2 bilinear upsampling is first adopted, followed by two residual blocks, each of which also has two 3 × 3 convolutions. Formally, the following operations are performed at *i*-*th* decoding scale:

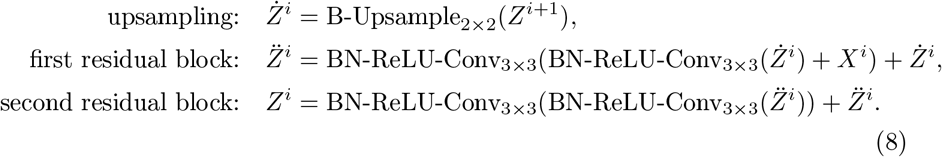

Note that the first residual block takes as input not only the previous feature map *Ż*^*i*^, but also the feature map *X*^*i*^ from the counterpart of the encoder. Such *skip connection*, which appends encoder features into the decoder features at the same resolutions, allows to retain the spatial details that are dropped during downsampling. Through the above encoding-decoding process, the GeneSegNet model generates an expressive yet full-resolution feature map, allowing for fine-grained analysis of the input image *I* and RNA location map *G*.

#### Network Output

The output of the decoder is a 32-channel feature map of full resolution, *i*.*e*., *Z*^0^ ∈ ℝ^32*×W×H*^. This feature map is separately fed into three different 1*×*1 convolutions. The first two convolutions are cascaded with sigmoid activation for respectively predicting the confidence map 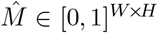 and the center map *Ĉ* ∈ [0, 1]^*W×H*^. The last convolution is to directly regress the two-channel offset map 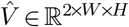. Formally, the final outputs of GeneSegNet are computed as:

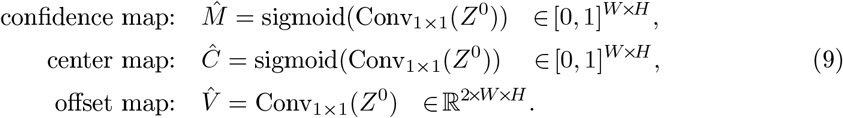

### Network Training

#### Image data processing

For both real datasets, we first split all images into training, validation, and testing sets. For NSCLC dataset, 59 raw images from two lung cancer samples are randomly split into 30 training images, 15 validation images, and 14 testing images. For the hippocampus dataset, 28 raw images were randomly split into 20 training images, 4 validation images, and 4 testing images. Since the raw images in both datasets are of very high resolution, it is infeasible to train and test deep learning models directly on the whole images due to the limitation of GPU memory. As a common practice, we randomly crop patch images with relatively small size of *W* × *H* from the raw training, validation, and test images, leading to thousands of patch images per dataset to be fed into GeneSegNet.

For the simulation dataset, since there is only one image in each simulation scenario, we randomly crop the image into smaller patch images with the same size of *W* × *H* and then split the patch images into 344 training, 100 validation, and 100 testing sets. Initial training labels were also cropped in the same way as the images. We empirically set *W* × *H* as 256 × 256, which is affordable for our GPUs.

#### Initial Training Label Generation

Before network training, we first derive the initial labels for the confidence map, center map, and offset map, from the instance segmentation annotations provided by the training datasets. Specifically, each training image *I* is associated with a set of *K* cell instance masks, *i*.*e*., 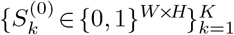. Note that the number of instances, *K*, varies across different images. For each pixel *p* = (*x, y*), the corresponding value in the *k*-*th* cell instance mask, *i*.*e*., 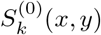, denotes whether *p* belongs to cell instance *k* (1) or not (0). Thus, the initial label *M*^(0)^∈{0, 1}^*W×H*^ for the confidence map is hence given as:

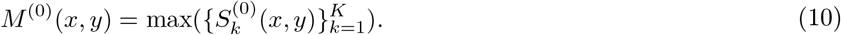

Therefore, *M*^(0)^(*x, y*) refers to if the pixel (*x, y*) is inside (1) or outside (0) of a certain cell instance.

The initial label *C*^(0)^∈{0, 1}^*W×H*^ for the center map is given as:

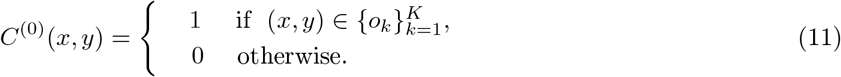

Here 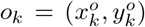 indicates the spatial coordinates of the center of *k*-*th* cell instance, computed by (median({*x*_*k*_}), median({*y*_*k*_})). {*x*_*k*_} and {*y*_*k*_} are the x- and y-axis coordinates of 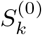. Thus *C*^(0)^ gathers all the instance centers.

The initial label *V*^(0)^∈ ℝ^2*×W ×H*^ for the offset map is given as:

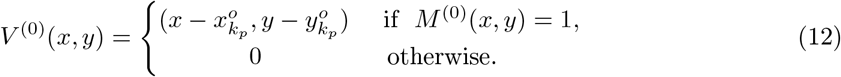

Here 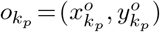 denotes the center position of the cell instance *k*_*p*_ that the cell pixel *p* belongs to. Thus, for each cell pixel *p* within a cell instance, *V*^(0)^(*p*) stores the offset from *p* = (*x, y*) to its corresponding center 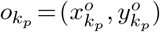.

#### Recursive Training Label optimization

As mentioned in §Methods Overview, we adopt a recursive training strategy (*cf*. Eq. 1) to mitigate the negative influence of noise in training labels. At each recursive training round *t* = {0, 1, 2, *…* }, after the converge of the network training (*cf*. Eq. 2), we use the updated network parameter *θ*^(*t*)^ to make predictions over the training data (***I, G***), *i*.*e*., 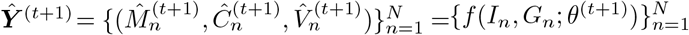, and a new set of *K*^*′*^ cell instance masks 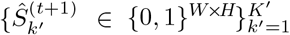 is further derived for each training sample (*I, G*). Then, for each mask 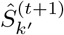, we apply the level-set algorithm [29] for further boundary refinement, which is given by:

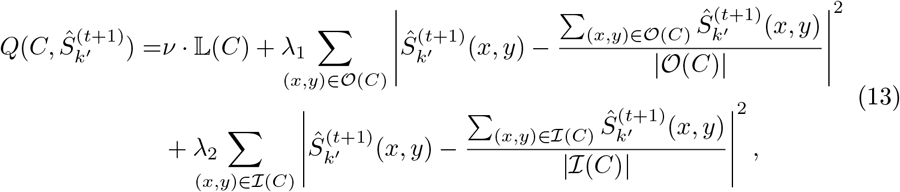

where *Q*(*·*) denotes the level set operation and |*·*| indicates the cardinality of a set. *C* represents the evolving contour of the *k*^*′*^-*th* cell instance mask 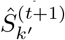 at (*t* + 1)-*th* recursive training round. 𝕃(*C*) denotes the length of the curve *C*. 𝒪(*C*) and ℐ(*C*) represent the regions outside and inside the contour *C. ν, λ*_1_ and *λ*_2_ are modulating parameters and are all set to be 1 in our study. Improved cell boundaries are obtained by minimizing the level set operation *Q*(*·*).

The initial contour of the cell instance mask 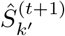 is defined implicitly via a Lipschitz function *φ*, by *C*_0_ = {(*x*_*c*_, *y*_*c*_) | *φ*(*x*_*c*_, *y*_*c*_) = 0}, which denotes the set of all contour points of the function *φ*(*z*_*c*_, *x*_*c*_, *y*_*c*_) = 0 at time *z*_*c*_ = 0, meaning zero level set. Therefore, the evolution of contour *C* can be carried out by gradually changing the value of the level set function *φ*(*·*) over time, which is equivalent to solving the following differential equation in the normal direction at each time:

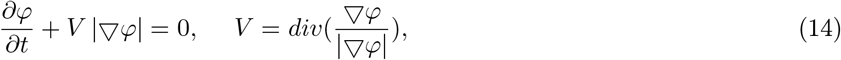

where *V* is the curvature of the function *φ*(*·*) passing through (*x*_*z*_, *y*_*z*_) [36, 29].

As the level-set algorithm takes into account the object’s geometric properties, such as curvature and gradient, it improves the convexity and elongation of the cell contour. The refined training labels 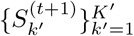 are used to create refined confidence map, center map, and offset map, *i*.*e*., 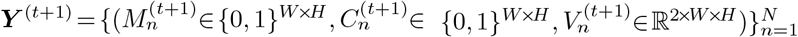, analogous to Eqs. 10, 11, and 12. The newly created training labels ***Y***^(*t*+1)^ are used for the further optimization of the network parameter.

#### Training Loss

Given current training labels 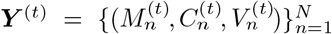 and network outputs 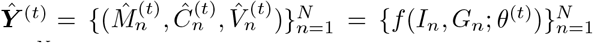 on the training data 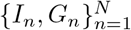, our training loss ℒ in Eq. 2 is defined as:

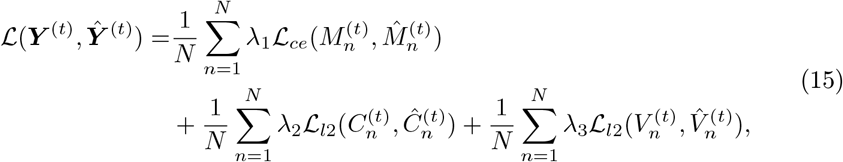

where ℒ_*ce*_ and ℒ_*l*2_ demonstrate the standard cross-entropy loss and L2 loss, respectively, which are applied in a position-wise manner; the coefficients are empirically set as *λ*_1_ = 1, *λ*_2_ = 1, and *λ*_3_ = 1, to balance the relative contributions among different training terms. Alg. 1 provides the pseudo code for the recursive network training strategy.

##### Algorithm 1 Recursive Training Procedure of GeneSegNet

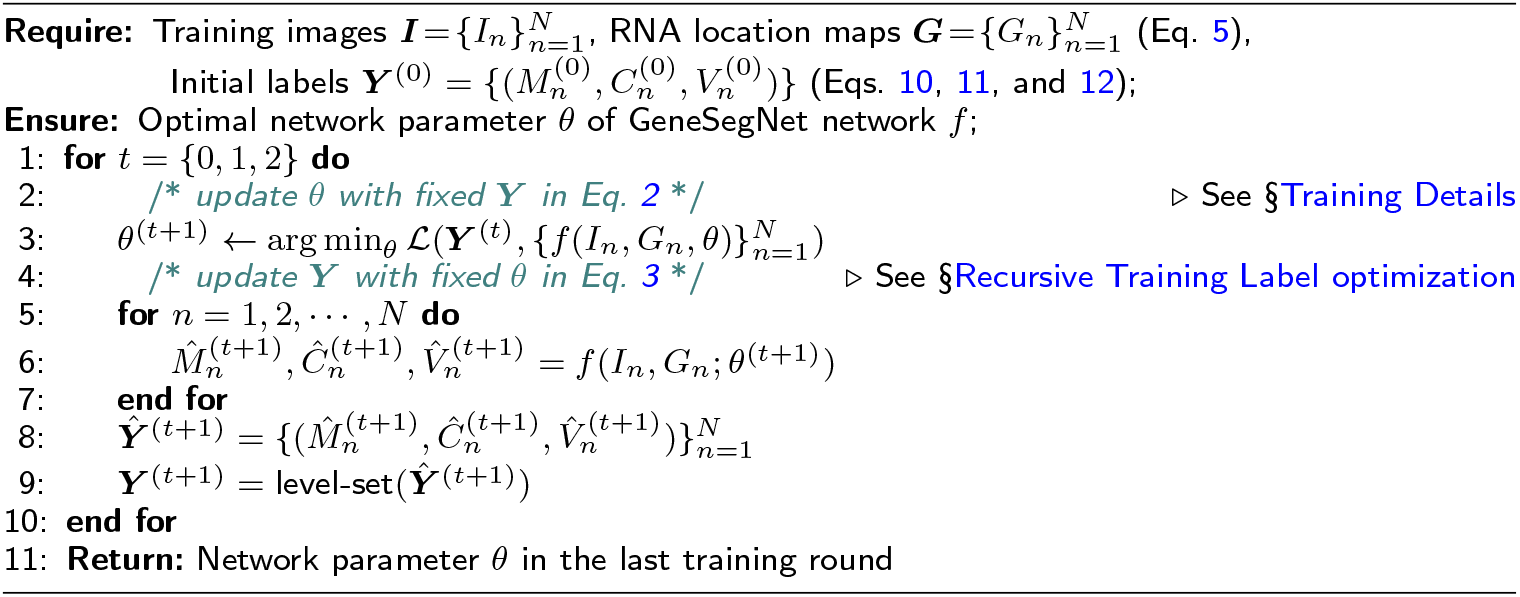

#### Training Details

GeneSegNet is trained in a recursive manner. In each recursive stage, the network *f* is trained for 500 epochs with a batch size of 8, a momentum of 0.9, and a weight decay of 1e-5. GeneSegNet was performed with different choices of learning rate (0.01, 0.001, and 0.0001), optimizer (Adam[37] and AdamW[28]), and variance *σ* (5, 7, and 9), and the set of parameters that leads to the highest cell calling score (3 pixels) was chosen. In an ablation study, we show that the performance of GeneSegNet is not hugely affected by different choices of parameters (Additional file 5-7: Table S3-S5). We use standard data augmentation techniques, including random scale jittering with a factor in [0.75, 1.25], random horizontal flipping, random cropping, and random rotation. By default, the recursive training procedure will stop after three iterations so that GeneSegNet finishes in reasonable time. In real practice, we also find that running three iterations is usually enough to reach a converged performance. However, users still have the freedom to set the number of iterative training rounds in real applications. GeneSegNet is implemented using PyTorch 1.11.0.

### Network Inference

Following the common practice [10] in this field, we divide the raw image into overlapping patches of the size of 256 × 256 pixels, with their corresponding RNA location maps. This allows GeneSegNet to handle images of arbitrary resolutions. These patches are overlapped both in vertical and horizontal directions by 50%, and fed into GeneSegNet individually for cell mask inference (*cf*. Eq. 4). Thus each pixel in the original image is processed four times. The final full-resolution segmentation is obtained through averaging the four predictions (in the coordinate system of the original image) for every pixel.

### Evaluation Metrics

#### Image Intersection Over Union (IoU) scores

Let 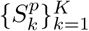 and 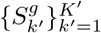 denote the predicted and ground truth cell instance masks, respectively. The IoU score for *k*-*th* cell instance is computed by:

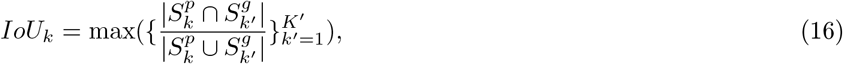

The overall image IoU score is the averaged cell-level image IoU scores across all cell instances.

#### Gene IoU scores

Let 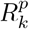 and 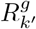 denote the sets of RNAs located inside the *k*-*th* predicted cell nstance 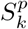 and the *k*^*′*^-*th* ground truth cell instance 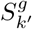, respectively. Here *k*^*′*^ = arg max, 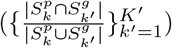. The gene IoU score for the *k*-*th* predicted cell instance is calculated as 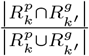. The overall gene IoU score is the averaged cell-level gene IoU scores across all cell instances.

#### Cell calling

Let 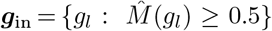 and 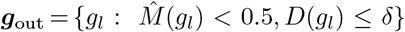 denote the sets of RNAs located inside or within a neighborhood outside predicted cell regions, respectively. *D*(*g*_*l*_) is the distance between *g*_*l*_ and the nearest pixel that belongs to a cell boundary. *δ* is a threshold and takes values of 3, 5, or 7. The cell calling metric is calculated as 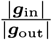, where | · |denotes the cardinality of a set. Compared to calculating cell calling scores separately within each cell and averaging cell calling scores across cells, the current approach is able to handle a situation where cells are crowded, RNAs from neighboring cells merge into a continuum, and differentiating RNAs within or outside cells is difficult. A higher cell calling means the method has a stronger ability to recruit RNAs inside cell boundaries.

#### Cell convexity

For each cell instance with boundary perimeter *r*_cell_, we define its convex hull as the smallest possible convex shape that completely contains the cell region and denote the perimeter of the convex hull as *r*_convex_. Then, the cell convexity of each cell is computed as 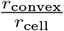.

#### Cell elongation

Let ***B*** be a set of two-dimensional points representing the boundary of a cell. We compute the covariance matrix of the ***B***, and then compute the two eigenvalues *e*_*a*_ and *e*_*b*_ of the covariance matrix, where *e*_*a*_ *> e*_*b*_. Cell elongation is computed as 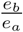.

#### Cell area

For each detected cell instance, its cell area is defined as the number of pixels it contains.

#### Reproducibility analysis

Cell clustering, differential gene analysis, and gene-gene correlation analysis were performed separately within each of the 28 mouse hippocampus samples and 2 NSCLC samples. For each hippocampus sample, segmented cells with positive expression in at least 5 genes were retained. For each NSCLC sample, segmented cells with positive expression in at least 30 genes were retained. Then for both hippocampus and NSCLC samples, the read count of each cell was divided by the total number of reads of that cell to produce normalized gene expression values. Genes with positive expression in at least 5% of cells were retained. Additional file 8: Table S6 includes lists of genes after filtering for different methods and for the two datasets. For cell clustering, normalized gene expression was used to group cells into 5 clusters by k-means clustering. The number of cells assigned to each cluster was divided by the total number of cells to produce proportions of cell clusters. The cell clusters were then used for differential gene analysis. For each gene and for each cell cluster, an averaged gene expression is calculated across all cells within that cell cluster. A fold change is calculated as the difference of averaged gene expression between a pair of cell clusters and for a gene. Fold changes across all pairs of clusters and all genes were concatenated into a single vector. For gene-gene correlations, Pearson correlation coefficient of normalized gene expression was calculated for each pair of genes across all cells.

For each pair of hippocampus samples from the same tissue (e.g., left hippocampus of the first mouse), a Pearson correlation is calculated between the two vectors of cell type proportions, differential gene analysis fold changes, or gene-gene correlations of all pairs of genes. The reproducibility is calculated as the median of correlations across all pairs of samples from the same tissue. For NSCLC dataset, the reproducibility is calculated as the Pearson correlation between the two vectors of cell type proportions, differential gene analysis fold changes, or gene-gene correlations of all pairs of genes between the two NSCLC samples. Additional file 9: Fig. S3 illustrates the process of calculating reproducibility for gene-gene correlations.

### Competing Methods

We compare GeneSegNet against five competing methods: Watershed algorithm [6, 7], Cellpose [10], JSTA [12], Baysor [11], and Baysor(prior) [11]. The implementation details are provided as follows.

#### Watershed Algorithm

For the hippocampus dataset, we directly obtained the segmentation results by the Watershed algorithm from the original publication [20]. For the simulation and NSCLC datasets, we implemented the Watershed algorithm using the nucleus images based on *scikit-image* [38] and *SciPy* [39] python packages. Specifically, we segment each image by (1) performing morphology erosion on the image to enable separations of cell instances that are adhesive together, (2) computing the distance map of the image and generating a set of markers as local maxima of the distance to the background, and (3) feeding the image and markers into the watershed algorithm in *scikit-image* for segmentation.

#### Cellpose Algorithm

For the simulation dataset, Cellpose [10] was trained on the simulated images with default settings and with labels generated by GeneSegNet model trained on hippocampus dataset. For the hippocampus dataset, Cellpose was trained using the DAPI nucleus staining images with default settings. For the NSCLC dataset, Cellpose segmentation with both nucleus and membrane images was obtained directly from the NanoString website. In addition, we also performed Cellpose with only the nucleus images using the default settings.

#### JSTA Algorithm

JSTA was performed with the default settings [12] for both real datasets. Spatial locations of cell centers and cell types of individual cells were directly downloaded from the original publication or NanoString website.

#### Baysor Algorithm

Baysor was performed with the default settings [11] for three datasets. In this mode, Baysor was performed without using imaging or prior segmentation information.

#### Baysor(prior) Algorithm

Baysor was performed with the default settings [11] for three datasets. In this mode, Baysor was performed with prior segmentation information. Specifically, for the simulation dataset, segmentation results from GeneSegNet model trained on the hippocampus dataset were used as prior segmentation information. For the NSCLC dataset, Cellpose segmentation results with both nucleus and membrane images were used as prior segmentation information. For the hippocampus dataset, Watershed segmentation results were used as prior segmentation information.

## Declarations

## Acknowledgements

We thank Drs. Kenneth D. Harris, Dimitris Nicoloutsopoulos, and Mats Nilsson for providing the mouse hippocampus dataset and its annotations. We also thank NanoString company for providing the NSCLC dataset.

## Funding

Z.J. was supported by the Whitehead Scholars Program at Duke University School of Medicine.

## Availability of data and materials

The simulation dataset [40] is available on figshare: https://doi.org/10.6084/m9.figshare.24012741 with License Apache-2.0, DOI 10.6084/m9.figshare.24012741.

The raw data of hippocampus [31] were downloaded from Figshare available from the original publication: https://doi.org/10.6084/m9.figshare.7150760 with License CC BY 4.0, DOI 10.6084/m9.figshare.7150760

The raw data of NSCLC datasets [32] were downloaded from NanoString website: https://nanostring.com/products/cosmx-spatial-molecular-imager/nsclc-ffpe-dataset/

GeneSegNet software package is available on GitHub [41]: https://github.com/BoomStarcuc/GeneSegNet with License MIT, on Code Ocean [42]: https://codeocean.com/capsule/9320020/tree/v2 with a configured environment, License MIT, or on Zenodo [43]: https://zenodo.org/record/8102672 with License MIT, DOI 10.5281/zenodo.8102672.

## Ethics approval and consent to participate

Not applicable.

## Competing interests

The authors declare that they have no competing interests

## Consent for publication

Not applicable.

## Authors’ contributions

Z.J. conceived the study. W.W., Y.W., D.L., T.Z., and Z.J. designed the algorithm and experiments. Y.W. performed all experiments with the aid of T.Z.. Y.W., W.H., and Z.J. analyzed the experiment results. All authors contributed to the writing of the manuscript.

## Additional Files

**Additional file 1: Fig. S1**. Model generalizability comparison of GeneSegNet and Cellpose.

**Additional file 2: Fig. S2**. Examples of ground truth cell segmentation in the NSCLC dataset.

**Additional file 3: Table S1**. Running time and memory usage of GeneSegNet.

**Additional file 4: Table S2**. Notation Table

**Additional file 5: Table S3**. The ablation study for the choice of hyper-parameters in the simulation dataset.

**Additional file 6: Table S4**. The ablation study for the choice of hyper-parameters in the NSCLC dataset.

**Additional file 7: Table S5**. The ablation study for the choice of hyper-parameters in the hippocampus dataset.

**Additional file 8: Table S6**. List of genes used in the reproducibility analysis for the NSCLC and hippocampus datasets.

**Additional file 9: Fig. S3**. Overview of the process calculating the reproducibility of gene-gene correlations.

## Notes

### Competing Interest Statement

The authors have declared no competing interest.

### Summary of Updates

We have added additional content in comparing across different cell segmentation methods.

## References

1. Crosetto, N., Bienko, M., Van Oudenaarden, A.: Spatially resolved transcriptomics and beyond. Nature Reviews Genetics 16(1), 57–66 (2015)

2. Chen, K.H., Boettiger, A.N., Moffitt, J.R., Wang, S., Zhuang, X.: Spatially resolved, highly multiplexed rna profiling in single cells. Science 348(6233), 6090 (2015)

3. Femino, A.M., Fay, F.S., Fogarty, K., Singer, R.H.: Visualization of single rna transcripts in situ. Science 280(5363), 585–590 (1998)

4. Eng, C.-H.L., Lawson, M., Zhu, Q., Dries, R., Koulena, N., Takei, Y., Yun, J., Cronin, C., Karp, C., Yuan, G.-C., et al.: Transcriptome-scale super-resolved imaging in tissues by rna seqfish+. Nature 568(7751), 235–239 (2019)

5. Wang, F., Flanagan, J., Su, N., Wang, L.-C., Bui, S., Nielson, A., Wu, X., Vo, H.-T., Ma, X.-J., Luo, Y.: Rnascope: a novel in situ rna analysis platform for formalin-fixed, paraffin-embedded tissues. The Journal of molecular diagnostics 14(1), 22–29 (2012)

6. Roerdink, J.B., Meijster, A.: The watershed transform: Definitions, algorithms and parallelization strategies. Fundamenta informaticae 41(1-2), 187–228 (2000)

7. Gamarra, M., Zurek, E., Escalante, H.J., Hurtado, L., San-Juan-Vergara, H.: Split and merge watershed: A two-step method for cell segmentation in fluorescence microscopy images. Biomedical signal processing and control 53, 101575 (2019)

8. Ronneberger, O., Fischer, P., Brox, T.: U-net: Convolutional networks for biomedical image segmentation. In: International Conference on Medical Image Computing and Computer-Assisted Intervention, pp. 234–241 (2015)

9. Van Valen, D.A., Kudo, T., Lane, K.M., Macklin, D.N., Quach, N.T., DeFelice, M.M., Maayan, I., Tanouchi, Y., Ashley, E.A., Covert, M.W.: Deep learning automates the quantitative analysis of individual cells in live-cell imaging experiments. PLoS computational biology 12(11), 1005177 (2016)

10. Stringer, C., Wang, T., Michaelos, M., Pachitariu, M.: Cellpose: a generalist algorithm for cellular segmentation. Nature Methods 18(1), 100–106 (2021)

11. Petukhov, V., Xu, R.J., Soldatov, R.A., Cadinu, P., Khodosevich, K., Moffitt, J.R., Kharchenko, P.V.: Cell segmentation in imaging-based spatial transcriptomics. Nature Biotechnology 40(3), 345–354 (2022)

12. Littman, R., Hemminger, Z., Foreman, R., Arneson, D., Zhang, G., Gómez-Pinilla, F., Yang, X., Wollman, R.: Joint cell segmentation and cell type annotation for spatial transcriptomics. Molecular Systems Biology 17(6), 10108 (2021)

13. Park, J., Choi, W., Tiesmeyer, S., Long, B., Borm, L.E., Garren, E., Nguyen, T.N., Tasic, B., Codeluppi, S., Graf, T., et al.: Cell segmentation-free inference of cell types from in situ transcriptomics data. Nature communications 12(1), 1–13 (2021)

14. Alom, M.Z., Hasan, M., Yakopcic, C., Taha, T.M., Asari, V.K.: Recurrent residual convolutional neural network based on u-net (r2u-net) for medical image segmentation. arXiv preprint arXiv:1802.06955 (2018)

15. Hofmanninger, J., Prayer, F., Pan, J., Röhrich, S., Prosch, H., Langs, G.: Automatic lung segmentation in routine imaging is primarily a data diversity problem, not a methodology problem. European Radiology Experimental 4(1), 1–13 (2020)

16. Zhang, C., Bengio, S., Hardt, M., Recht, B., Vinyals, O.: Understanding deep learning (still) requires rethinking generalization. Communications of the ACM 64(3), 107–115 (2021)

17. Reed, S., Lee, H., Anguelov, D., Szegedy, C., Erhan, D., Rabinovich, A.: Training deep neural networks on noisy labels with bootstrapping. arXiv preprint arXiv:1412.6596 (2014)

18. Arpit, D., Jastrzębski, S., Ballas, N., Krueger, D., Bengio, E., Kanwal, M.S., Maharaj, T., Fischer, A., Courville, A., Bengio, Y., et al.: A closer look at memorization in deep networks. In: International Conference on Machine Learning, pp. 233–242 (2017)

19. He, S., Bhatt, R., Brown, C., Brown, E.A., Buhr, D.L., Chantranuvatana, K., Danaher, P., Dunaway, D., Garrison, R.G., Geiss, G., et al.: High-plex imaging of rna and proteins at subcellular resolution in fixed tissue by spatial molecular imaging. Nature Biotechnology 40(12), 1794–1806 (2022)

20. Qian, X., Harris, K.D., Hauling, T., Nicoloutsopoulos, D., Muñoz-Manchado, A.B., Skene, N., Hjerling-Leffler, J., Nilsson, M.: Probabilistic cell typing enables fine mapping of closely related cell types in situ. Nature Methods 17(1), 101–106 (2020)

21. Girshick, R.: Fast r-cnn. In: Proceedings of the IEEE International Conference on Computer Vision, pp. 1440–1448 (2015)

22. Redmon, J., Divvala, S., Girshick, R., Farhadi, A.: You only look once: Unified, real-time object detection. In: Proceedings of the IEEE Conference on Computer Vision and Pattern Recognition, pp. 779–788 (2016)

23. He, K., Gkioxari, G., Dollár, P., Girshick, R.: Mask r-cnn. In: Proceedings of the IEEE International Conference on Computer Vision, pp. 2961–2969 (2017)

24. Schmidt, U., Weigert, M., Broaddus, C., Myers, G.: Cell detection with star-convex polygons. In: Medical Image Computing and Computer Assisted Intervention–MICCAI 2018: 21st International Conference, Granada, Spain, September 16-20, 2018, Proceedings, Part II 11, pp. 265–273 (2018). Springer

25. Langfelder, P., Horvath, S.: Wgcna: an r package for weighted correlation network analysis. BMC bioinformatics 9(1), 1–13 (2008)

26. Junttila, S., Smolander, J., Elo, L.L.: Benchmarking methods for detecting differential states between conditions from multi-subject single-cell rna-seq data. Briefings in bioinformatics 23(5), 286 (2022)

27. Kang, Y., Thieffry, D., Cantini, L.: Evaluating the reproducibility of single-cell gene regulatory network inference algorithms. Frontiers in genetics 12, 362 (2021)

28. Loshchilov, I., Hutter, F.: Decoupled weight decay regularization. arXiv preprint arXiv:1711.05101 (2017)

29. Osher, S., Sethian, J.A.: Fronts propagating with curvature-dependent speed: Algorithms based on hamilton-jacobi formulations. Journal of Computational Physics 79(1), 12–49 (1988)

30. Li, C., Xu, C., Gui, C., Fox, M.D.: Distance regularized level set evolution and its application to image segmentation. IEEE Transactions on Image Processing 19(12), 3243–3254 (2010)

31. Qian, X., Harris, K.D., Hauling, T., Nicoloutsopoulos, D., Muñoz-Manchado, A.B., Skene, N., Hjerling-Leffler, J., Nilsson, M.: Probabilistic cell typing enables fine mapping of closely related cell types in situ. Mouse hippocampus CA1 region (hippocampus) dataset (2019). doi:10.6084/m9.figshare.7150760. https://su.figshare.com/articles/dataset/pciSeq_files_in_csv_format/10318610

32. He, S., Bhatt, R., Brown, C., Brown, E.A., Buhr, D.L., Chantranuvatana, K., Danaher, P., Dunaway, D., Garrison, R.G., Geiss, G., et al.: High-plex imaging of rna and proteins at subcellular resolution in fixed tissue by spatial molecular imaging. Human non-small cell lung cancer (NSCLC) dataset (2022). https://nanostring.com/products/cosmx-spatial-molecular-imager/nsclc-ffpe-dataset/

33. Svoboda, D., Kozubek, M., Stejskal, S.: Generation of digital phantoms of cell nuclei and simulation of image formation in 3d image cytometry. Cytometry Part A: The Journal of the International Society for Advancement of Cytometry 75(6), 494–509 (2009)

34. Cao, Z., Simon, T., Wei, S.-E., Sheikh, Y.: Realtime multi-person 2d pose estimation using part affinity fields. In: Proceedings of the IEEE Conference on Computer Vision and Pattern Recognition, pp. 7291–7299 (2017)

35. He, K., Zhang, X., Ren, S., Sun, J.: Deep residual learning for image recognition. In: Proceedings of the IEEE Conference on Computer Vision and Pattern Recognition, pp. 770–778 (2016)

36. Chan, T.F., Vese, L.A.: Active contours without edges. IEEE Transactions on Image Processing 10(2), 266–277 (2001)

37. Kingma, D.P., Ba, J.: Adam: A method for stochastic optimization. arXiv preprint arXiv:1412.6980 (2014)

38. van der Walt, S., Schönberger, J.L., Nunez-Iglesias, J., Boulogne, F., Warner, J.D., Yager, N., Gouillart, E., Yu, T., the scikit-image contributors: scikit-image: image processing in Python. PeerJ 2, 453 (2014)

39. Virtanen, P., Gommers, R., Oliphant, T.E., Haberland, M., Reddy, T., Cournapeau, D., Burovski, E., Peterson, P., Weckesser, W., Bright, J., van der Walt, S.J., Brett, M., Wilson, J., Millman, K.J., Mayorov, N., Nelson, A.R.J., Jones, E., Kern, R., Larson, E., Carey, C.J., Polat, I., Feng, Y., Moore, E.W., VanderPlas, J., Laxalde, D., Perktold, J., Cimrman, R., Henriksen, I., Quintero, E.A., Harris, C.R., Archibald, A.M., Ribeiro, A.H., Pedregosa, F., van Mulbregt, P., SciPy 1.0 Contributors: SciPy 1.0: Fundamental Algorithms for Scientific Computing in Python. Nature Methods 17, 261–272 (2020)

40. Wang, Y., Wang, W., Liu, D., Hou, W., Zhou, T., Ji, Z.: GeneSegNet: a deep learning framework for cell segmentation by integrating gene expression and imaging. Simulation dataset (2023). doi:10.6084/m9.figshare.24012741. 10.6084/m9.figshare.24012741

41. Wang, Y., Wang, W., Liu, D., Hou, W., Zhou, T., Ji, Z.: GeneSegNet: a deep learning framework for cell segmentation by integrating gene expression and imaging. Github (2023). https://github.com/BoomStarcuc/GeneSegNet

42. Wang, Y., Wang, W., Liu, D., Hou, W., Zhou, T., Ji, Z.: GeneSegNet: a deep learning framework for cell segmentation by integrating gene expression and imaging. Code Ocean (2023). doi:10.24433/CO.4787636.v2. https://codeocean.com/capsule/9320020/tree/v2

43. Wang, Y., Wang, W., Liu, D., Hou, W., Zhou, T., Ji, Z.: GeneSegNet: a deep learning framework for cell segmentation by integrating gene expression and imaging. Zenodo (2023). doi:10.5281/zenodo.8102672. https://zenodo.org/record/8102672

