## Supplementary Materials for "GeneSegNet: a deep learning framework for cell segmentation by integrating gene expression and imaging"

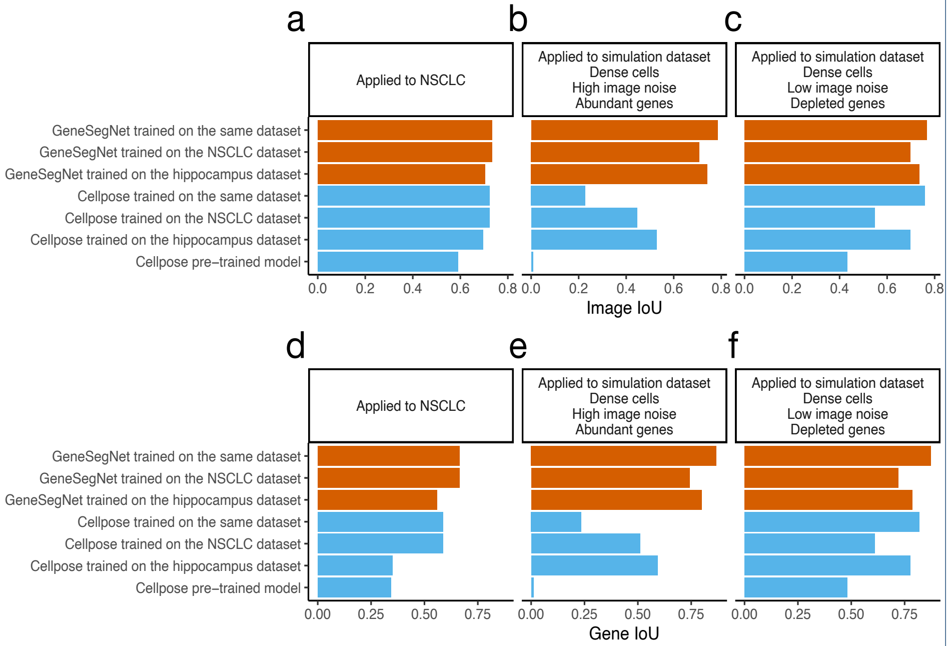


**Fig. S1:** Image IoU scores (**a-c**) and Gene IoU scores (**d-f**) when GeneSegNet and Cellpose were trained on the same or different datasets as the dataset being applied. **a,d**: Models applied to NSCLC dataset. **b,e**: Models applied to simulation dataset with densely distributed cells, high image noise, and abundant amount of genes. **c,f**: Models applied to simulation dataset with densely distributed cells, low image noise, and depleted amount of genes.

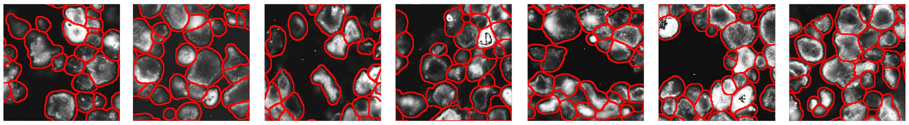

**Fig. S2:** Examples of ground truth cell segmentation derived by Cellpose using both nuclear and membrane markers in the NSCLC dataset. The detected cell boundaries are marked in red, shown on membrane images as background.

**Table S1:** Running time and memory usage of GeneSegNet.

| Dataset | GeneSegNet Stage | Runtime | Memory Usage | Configuration |
|---|---|---|---|---|
| Simulation (one scenario) | Training | 330 Min | 8GB GPU Memory | Image Size = 256 × 256, Epoch = 500, Batch Size = 8, Training and Validation Samples = 344 + 100 |
| NSCLC | | 720 Min | | Image Size = 256 × 256, Epoch = 500, Batch Size = 8, Training and Validation Samples = 961 + 290 |
| Hippocampus | | 1140 Min | | Image Size = 256 × 256, Epoch = 500, Batch Size = 8, Training and Validation Samples = 2098 + 420 |
| Simulation (one scenario) | Inference | 20.0 Min | 20GB CPU Memory | Image Size = 7248 × 3624 |
| NSCLC | | 15.0 Min | 10GB CPU Memory | Image Size = 5472 × 3648 |
| Hippocampus | | 20.0 Min | 30GB CPU Memory | Image Size = 6130 × 5548 |

**Table S2:** Notation Table

| Notation | Description |
|---|---|
| $I$ | input (patch) image of $W \times H$ resolution, $I \in [0,1]^{W\times H}$ |
| $g_l$ | coordinate of the $l$-th RNA in the lattice of $I$, $g_l \in \{1,\cdots,W\}\times\{1,\cdots,H\}$ |
| $G$ | input RNA location map, $G \in [0,1]^{W\times H}$ |
| $\boldsymbol{I}$ | set of a total of $N$ training (patch) images, $\boldsymbol{I} = \{I_n\}_{n=1}^{N}$ |
| $\boldsymbol{g}$ | set of coordinates of a total of $L$ RNAs in image $I$, $\boldsymbol{g} = \{g_l\}_{l=1}^{L}$ |
| $\boldsymbol{G}$ | set of a total of $N$ RNA location maps of the training images $\boldsymbol{I}$, $\boldsymbol{G} = \{G_n\}_{n=1}^{N}$ |
| $f(;\theta)$ | neural network of GeneSegNet parameterized by $\theta$ |
| $\hat{M}/M$ | network estimation/training label for the confidence map, $\hat{M} \in [0,1]^{W\times H}$, $M \in \{0,1\}^{W\times H}$ |
| $\hat{C}/C$ | network estimation/training label for the center map, $\hat{C} \in [0,1]^{W\times H}$, $C \in \{0,1\}^{W\times H}$ |
| $\hat{V}/V$ | network estimation/training label for the offset map, $\hat{V}, V \in \mathbb{R}^{2\times W\times H}$ |
| $\hat{\boldsymbol{Y}}/\boldsymbol{Y}$ | set of network outputs/training labels for the training images $\boldsymbol{I}$, $\hat{\boldsymbol{Y}} = \{(\hat{M}_n, \hat{C}_n, \hat{V}_n)\}_{n=1}^{N}$, $\boldsymbol{Y} = \{(M_n, C_n, V_n)\}_{n=1}^{N}$ |
| $\hat{S}_k/S_k$ | network estimation/training label for the mask of $k$-*th* cell instance in image $I$, $\hat{S}_k, S_k \in \{0,1\}^{W\times H}$ |
| $\hat{o}_k/o_k$ | network estimation/training label for center coordinate of $k$-*th* cell instance in the lattice of $I$ |
| $\mathcal{L}$, $\mathcal{L}_{ce}$, $\mathcal{L}_{l2}$ | loss functions used for network optimization |

**Table S3:** The ablation study for the choice of hyper-parameters (optimizer, learning rate, variance $\sigma$) in simulation dataset with image and gene IoU scores. The performances corresponding to the chosen parameters are highlighted in bold.

| Optimizer | Learning Rate | Dataset | Image IoU | | | | | Gene IoU | | | | |
|---|---|---|---|---|---|---|---|---|---|---|---|---|
| | | | Variance $\sigma$ | | | | | Variance $\sigma$ | | | | |
| | | | 3 | 5 | 7 | 9 | 11 | 3 | 5 | 7 | 9 | 11 |
| SGD | 0.0001 | Dense cells High image noise Abundant genes | 0.685 | 0.682 | 0.731 | 0.761 | 0.746 | 0.734 | 0.731 | 0.831 | 0.854 | 0.841 |
| | 0.001 | | 0.735 | 0.74 | 0.755 | 0.769 | 0.75 | 0.817 | 0.826 | 0.843 | 0.858 | 0.838 |
| | 0.01 | | 0.726 | 0.728 | 0.688 | 0.737 | 0.724 | 0.812 | 0.819 | 0.77 | 0.844 | 0.83 |
| | 0.1 | | 0.703 | 0.794 | 0.702 | 0.687 | 0.695 | 0.772 | 0.764 | 0.781 | 0.753 | 0.778 |
| Adam | 0.0001 | | 0.668 | 0.721 | 0.740 | 0.774 | 0.688 | 0.712 | 0.814 | 0.836 | 0.86 | 0.751 |
| | 0.001 | | 0.712 | 0.732 | 0.774 | 0.783 | 0.74 | 0.802 | 0.821 | 0.853 | 0.877 | 0.836 |
| | 0.01 | | 0.693 | 0.707 | 0.752 | 0.767 | 0.764 | 0.760 | 0.781 | 0.841 | 0.858 | 0.855 |
| | 0.1 | | 0.665 | 0.679 | 0.716 | 0.706 | 0.731 | 0.710 | 0.727 | 0.824 | 0.79 | 0.832 |
| AdamW | 0.0001 | | 0.731 | 0.719 | 0.763 | **0.779** | 0.737 | 0.812 | 0.813 | 0.85 | **0.865** | 0.835 |
| | 0.001 | | 0.719 | 0.753 | 0.771 | 0.788 | 0.755 | 0.808 | 0.835 | 0.851 | 0.883 | 0.847 |
| | 0.01 | | 0.679 | 0.747 | 0.748 | 0.733 | 0.77 | 0.723 | 0.83 | 0.838 | 0.838 | 0.854 |
| | 0.1 | | 0.697 | 0.69 | 0.714 | 0.713 | 0.716 | 0.763 | 0.745 | 0.818 | 0.824 | 0.822 |

| Optimizer | Learning Rate | Dataset | Image IoU | | | | | Gene IoU | | | | |
|---|---|---|---|---|---|---|---|---|---|---|---|---|
| | | | Variance $\sigma$ | | | | | Variance $\sigma$ | | | | |
| | | | 3 | 5 | 7 | 9 | 11 | 3 | 5 | 7 | 9 | 11 |
| SGD | 0.0001 | Dense cells Low image noise Depleted genes | 0.644 | 0.677 | 0.758 | 0.699 | 0.673 | 0.766 | 0.808 | 0.863 | 0.824 | 0.82 |
| | 0.001 | | 0.685 | 0.741 | 0.762 | 0.738 | 0.700 | 0.811 | 0.844 | 0.868 | 0.84 | 0.841 |
| | 0.01 | | 0.668 | 0.705 | 0.741 | 0.719 | 0.726 | 0.796 | 0.828 | 0.849 | 0.831 | 0.85 |
| | 0.1 | | 0.638 | 0.644 | 0.661 | 0.653 | 0.658 | 0.759 | 0.766 | 0.802 | 0.781 | 0.803 |
| Adam | 0.0001 | | 0.653 | 0.650 | 0.745 | 0.733 | 0.690 | 0.768 | 0.779 | 0.85 | 0.838 | 0.829 |
| | 0.001 | | 0.710 | 0.744 | **0.764** | 0.752 | 0.750 | 0.823 | 0.847 | **0.871** | 0.851 | 0.862 |
| | 0.01 | | 0.688 | 0.716 | 0.695 | 0.674 | 0.734 | 0.808 | 0.833 | 0.83 | 0.805 | 0.854 |
| | 0.1 | | 0.629 | 0.669 | 0.656 | 0.712 | 0.713 | 0.754 | 0.803 | 0.788 | 0.829 | 0.845 |
| AdamW | 0.0001 | | 0.681 | 0.658 | 0.754 | 0.718 | 0.699 | 0.8 | 0.792 | 0.858 | 0.831 | 0.844 |
| | 0.001 | | 0.708 | 0.727 | 0.759 | 0.761 | 0.738 | 0.822 | 0.837 | 0.86 | 0.861 | 0.856 |
| | 0.01 | | 0.722 | 0.739 | 0.774 | 0.748 | 0.746 | 0.828 | 0.842 | 0.884 | 0.844 | 0.86 |
| | 0.1 | | 0.660 | 0.692 | 0.702 | 0.730 | 0.721 | 0.79 | 0.819 | 0.834 | 0.836 | 0.846 |

**Table S4:** The ablation study for the choice of hyper-parameters (optimizer, learning rate, variance $\sigma$) in NSCLC dataset with image and gene IoU scores. The performances corresponding to the chosen parameters are highlighted in bold.

| Optimizer | Learning Rate | Dataset | Image IoU | | | | | Gene IoU | | | | |
|---|---|---|---|---|---|---|---|---|---|---|---|---|
| | | | Variance $\sigma$ | | | | | Variance $\sigma$ | | | | |
| | | | 3 | 5 | 7 | 9 | 11 | 3 | 5 | 7 | 9 | 11 |
| SGD | 0.0001 | NSCLC | 0.688 | 0.683 | 0.676 | 0.692 | 0.67 | 0.759 | 0.752 | 0.747 | 0.766 | 0.74 |
| | 0.001 | | 0.708 | 0.696 | 0.672 | 0.702 | 0.692 | 0.787 | 0.766 | 0.744 | 0.779 | 0.765 |
| | 0.01 | | 0.673 | 0.634 | 0.685 | 0.714 | 0.701 | 0.746 | 0.699 | 0.756 | 0.8 | 0.781 |
| | 0.1 | | 0.659 | 0.647 | 0.648 | 0.676 | 0.654 | 0.718 | 0.71 | 0.712 | 0.748 | 0.716 |
| Adam | 0.0001 | | 0.711 | 0.663 | 0.697 | 0.715 | 0.707 | 0.797 | 0.725 | 0.768 | 0.8 | 0.786 |
| | 0.001 | | 0.713 | 0.673 | 0.708 | 0.72 | 0.714 | 0.798 | 0.745 | 0.787 | 0.804 | 0.798 |
| | 0.01 | | 0.687 | 0.69 | 0.657 | 0.691 | 0.699 | 0.757 | 0.764 | 0.718 | 0.764 | 0.773 |
| | 0.1 | | 0.668 | 0.629 | 0.633 | 0.684 | 0.649 | 0.727 | 0.693 | 0.699 | 0.756 | 0.712 |
| AdamW | 0.0001 | | 0.721 | 0.685 | 0.691 | 0.728 | 0.709 | 0.804 | 0.755 | 0.764 | 0.809 | 0.787 |
| | 0.001 | | 0.719 | 0.689 | 0.686 | **0.734** | 0.711 | 0.801 | 0.76 | 0.756 | **0.817** | 0.796 |
| | 0.01 | | 0.692 | 0.658 | 0.668 | 0.706 | 0.684 | 0.765 | 0.72 | 0.727 | 0.785 | 0.754 |
| | 0.1 | | 0.678 | 0.641 | 0.664 | 0.689 | 0.662 | 0.75 | 0.704 | 0.725 | 0.761 | 0.721 |

**Table S5:** The ablation study for the choice of hyper-parameters (optimizer, learning rate, variance $\sigma$) in the hippocampus dataset. The performances corresponding to the chosen parameters are highlighted in bold.

| Optimizer | Learning Rate | Dataset | Variance $\sigma$ | | | | | | | | | |
|---|---|---|---|---|---|---|---|---|---|---|---|---|
| | | | 3 | | | | | 5 | | | | |
| | | | Cell Calling | | | Cell Area (pixel) | | Cell Calling | | | Cell Area (pixel) | |
| | | | 3 pixel | 5 pixel | 7 pixel | Avg Area | Avg Radius | 3 pixel | 5 pixel | 7 pixel | Avg Area | Avg Radius |
| SGD | 0.0001 | Hippocampus | 37.93 | 18.02 | 10.09 | 1119.47 | 18.88 | 34.33 | 13.89 | 7.02 | 1057.20 | 18.35 |
| | 0.001 | | 39.84 | 19.58 | 11.63 | 1142.15 | 19.07 | 36.95 | 16.91 | 9.32 | 1109.14 | 18.79 |
| | 0.01 | | 44.20 | 21.90 | 14.80 | 1199.14 | 19.54 | 36.50 | 16.65 | 8.99 | 1103.48 | 18.75 |
| | 0.1 | | 36.89 | 16.72 | 9.05 | 1105.58 | 18.76 | 35.19 | 15.27 | 7.90 | 1064.32 | 18.41 |
| Adam | 0.0001 | | 36.37 | 16.26 | 8.91 | 1100.74 | 18.72 | 35.56 | 15.76 | 8.18 | 1077.49 | 18.52 |
| | 0.001 | | 42.24 | 21.90 | 14.05 | 1173.87 | 19.34 | 38.23 | 18.27 | 10.70 | 1124.86 | 18.93 |
| | 0.01 | | 41.66 | 20.37 | 13.43 | 1160.96 | 19.23 | 37.77 | 17.83 | 9.95 | 1117.37 | 18.86 |
| | 0.1 | | 39.17 | 18.94 | 11.21 | 1128.84 | 18.96 | 34.85 | 14.30 | 7.44 | 1058.19 | 18.36 |
| AdamW | 0.0001 | | 37.55 | 17.18 | 9.40 | 1110.21 | 18.80 | 36.04 | 16.18 | 8,76 | 1083.26 | 18.57 |
| | 0.001 | | 40.45 | 20.22 | 12.70 | 1159.33 | 19.21 | 39.68 | 20.31 | 13.44 | 1153.11 | 19.16 |
| | 0.01 | | 42.75 | 22.62 | 14.77 | 1175.26 | 19.35 | 38.88 | 18.54 | 11.62 | 1126.93 | 18.94 |
| | 0.1 | | 38.51 | 18.47 | 10.62 | 1126.03 | 18.94 | 35.02 | 14.96 | 7.81 | 1062.27 | 18.39 |

| Optimizer | Learning Rate | Dataset | Variance $\sigma$ | | | | | | | | | |
|---|---|---|---|---|---|---|---|---|---|---|---|---|
| | | | 7 | | | | | 9 | | | | |
| | | | Cell Calling | | | Cell Area (pixel) | | Cell Calling | | | Cell Area (pixel) | |
| | | | 3 pixel | 5 pixel | 7 pixel | Avg Area | Avg Radius | 3 pixel | 5 pixel | 7 pixel | Avg Area | Avg Radius |
| SGD | 0.0001 | Hippocampus | 46.30 | 23.76 | 15.68 | 1297.43 | 20.33 | 38.26 | 18.18 | 10.24 | 1121.01 | 18.90 |
| | 0.001 | | 47.63 | 24.20 | 16.01 | 1300.94 | 20.35 | 40.02 | 19.92 | 11.86 | 1155.11 | 19.18 |
| | 0.01 | | 46.82 | 24.03 | 15.70 | 1298.92 | 20.34 | 42.01 | 21.88 | 13.05 | 1168.47 | 19.29 |
| | 0.1 | | 45.37 | 23.62 | 15.64 | 1295.36 | 20.31 | 37.74 | 17.56 | 9.61 | 1116.50 | 18.86 |
| Adam | 0.0001 | | 47.18 | 24.11 | 15.79 | 1299.90 | 20.35 | 40.38 | 20.11 | 12.43 | 1158.04 | 19.20 |
| | 0.001 | | 47.28 | 24.13 | 15.87 | 1300.24 | 20.35 | 44.41 | 23.52 | 15.64 | 1292.87 | 20.29 |
| | 0.01 | | 48.33 | 24.92 | 16.02 | 1311.58 | 20.44 | 42.12 | 22.09 | 13.78 | 1172.22 | 19.32 |
| | 0.1 | | 46.67 | 23.94 | 15.68 | 1298.58 | 20.34 | 38.87 | 18.69 | 10.77 | 1125.56 | 18.93 |
| AdamW | 0.0001 | | 49.87 | 25.17 | 16.91 | 1352.35 | 20.75 | 41.64 | 20.41 | 12.88 | 1160.30 | 19.22 |
| | 0.001 | | **50.76** | **25.24** | **17.88** | **1364.79** | **20.85** | 43.79 | 23.07 | 14.92 | 1188.68 | 19.46 |
| | 0.01 | | 48.62 | 24.98 | 16.33 | 1312.21 | 20.44 | 42.81 | 22.78 | 14.37 | 1176.14 | 19.35 |
| | 0.1 | | 48.08 | 24.36 | 16.01 | 1303.32 | 20.37 | 39.47 | 19.21 | 11.12 | 1130.29 | 18.97 |

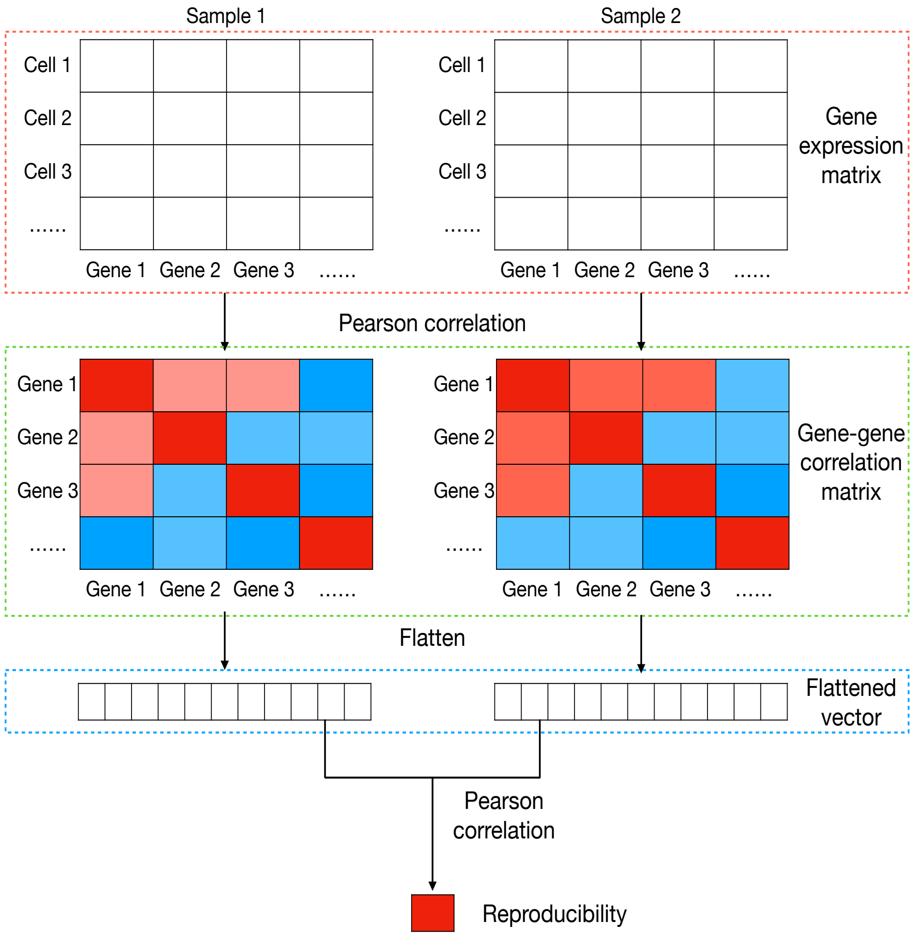


**Fig. S3:** Overview of the process calculating the reproducibility of gene-gene correlations. Red and blue colors in the matrices indicate positive and negative values, respectively.
